# A Systematic Comparison of Differential Analysis Methods for CyTOF Data

**DOI:** 10.1101/2021.08.09.455609

**Authors:** Lis Arend, Judith Bernett, Quirin Manz, Melissa Klug, Olga Lazareva, Jan Baumbach, Dario Bongiovanni, Markus List

## Abstract

Cytometry techniques are widely used to discover cellular characteristics at single-cell resolution. Many data analysis methods for cytometry data focus solely on identifying subpopulations via clustering and testing for differential cell abundance. For differential expression analysis of markers between conditions, only few tools exist. These tools either reduce the data distribution to medians, discarding valuable information, or have underlying assumptions that may not hold for all expression patterns.

Here, we systematically evaluated existing and novel approaches for differential expression analysis on real and simulated CyTOF data. We found that methods using median marker expressions compute fast and reliable results when the data is not strongly zero-inflated. Methods using all data detect changes in strongly zero-inflated markers, but partially suffer from overprediction or cannot handle big datasets. We present a new method, CyEMD, based on calculating the Earth Mover’s Distance between expression distributions that can handle strong zero-inflation without being too sensitive.

Additionally, we developed CYANUS, a user-friendly R Shiny App allowing the user to analyze cytometry data with state-of-the-art tools, including well-performing methods from our comparison. A public web interface is available at https://exbio.wzw.tum.de/cyanus/.

## Introduction

In conventional flow cytometry, single cells are passed through one or multiple lasers while being suspended in a liquid stream. Antibodies are labeled with fluorescent dyes and lasers produce both scattered and fluorescent light signals that are read by detectors (***McKinnon*** (***2018***)). This enables the analysis, identification, and classification of cell populations which is required in multiple disciplines such as immunology, cancer biology, and virology. However, flow cytometry has some severe limitations restricting its utility. The number of parameters analyzed at one time is limited due to the overlap of the light emissions, the rupture of the stains, and the requirement of large cell numbers. (***Gadalla et al.*** (***2019***))

Recently, high-dimensional time-of-flight mass cytometry (CyTOF) has emerged with the ability to identify more than 40 parameters simultaneously. Its advantage over flow cytometry is that antibodies are labeled with metal isotopes instead of fluorophores, allowing scientists to analyze more antibodies in a single run while needing fewer cells per experiment. Traditional flow cytometry would require multiple tubes with different antibody panels to cover the same number of markers (***Gadalla et al.*** (***2019***)). Consequently, CyTOF experiments are a powerful tool to unveil new cell subtypes, functions, and biomarkers in many fields, e.g. the discovery of disease-associated immunologic changes in cancer.

Cytometry experiments rely on a panel of antibodies that are associated with a specific experimental condition or phenotype of interest. Usually, the analysis of cytometry data starts by clustering cells into cell subpopulations, followed by a differential expression analysis between and within cell types (***Nowicka et al.*** (***2019***)). Several methods have been developed for testing clusters representing cell populations for differential abundance (DA) between conditions (***Bruggner et al.*** (*2014*), ***Arvaniti and Claassen*** (***2017***), ***Weber et al.*** (***2019***)). However, many experiments aim to detect differential states (DS), i.e. differential expression of markers between conditions and within cell populations (see Figure 1A).

**Figure 1.**
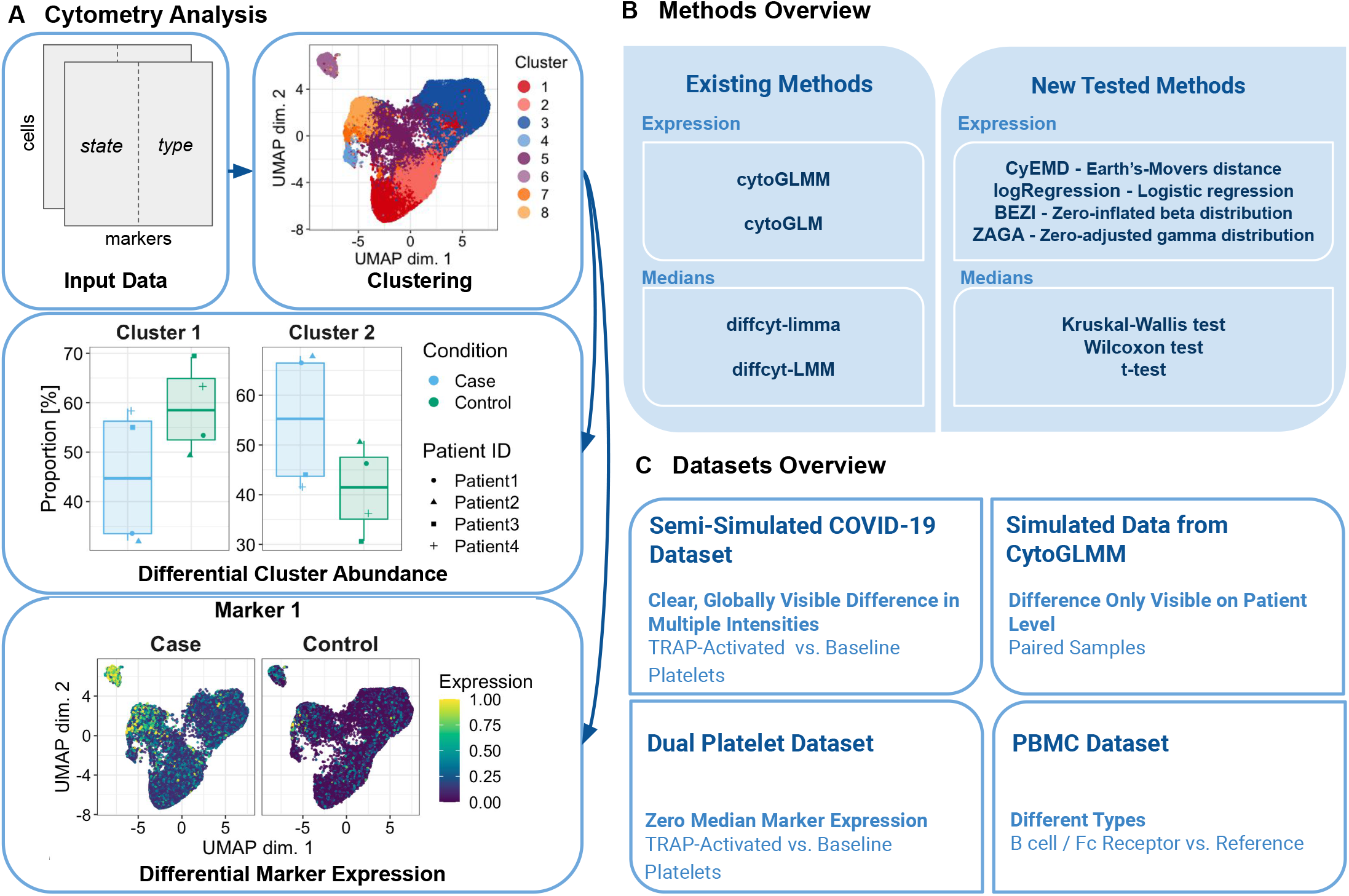
**(A)** Schematic overview of a differential analysis workflow for cytometry data. In a cytometry experiment, the abundance of state (condition) and type (lineage) markers are measured for each cell. Usually, cells are clustered using type markers to identify cell subpopulations. When differential cluster abundance is analysed, the proportion of cell types between conditions is compared (e.g. condition 2 stimulates the production of cell subpopulation 1). When differential marker expression is analysed, marker expression is compared between conditions within each cluster (e.g. Marker 1 is more highly expressed in condition 1 in clusters 6 and 8). **(B)** Overview of the methods compared in this study. **(C)** Overview of the datasets used in this study. One simulated, one semi-simulated, and two real CyTOF datasets were used to evaluate the methods.

Diffcyt (***Weber et al.*** (***2019***)) presents two methods for differential expression detection, a linear mixed effect model (LMM) and an adaptation of limma (***Ritchie et al*.** (***2015***)). For both approaches, the data is reduced to median marker expressions per sample and per cluster when comparing conditions. Another recently developed method is CytoGLMM (***Seiler et al.*** (***2021***)) which introduces two multiple regression strategies for finding differential proteins in mass cytometry data: a bootstrapped generalized linear model and a generalized linear mixed model allowing for random and fixed effects.

Methods that rely solely on median marker expression and do not take other distribution characteristics into account might be oblivious to certain marker expression patterns. At the same time, the comparison of hundreds of thousands of cells per patient is computationally and statistically tedious. In this study, we provide a clear overview of existing and novel approaches and compare them in different scenarios, highlighting their strengths and weaknesses. As novel approaches, we implemented three statistical tests relying on the medians, a logistic regression using all expression data, two techniques modeling the expression distributions and a method using the Earth Mover’s distance (see Figure 1B). A similar approach to the latter, SigEMD, has recently been introduced by ***Wang and Nabavi*** (***2018***) for single-cell RNA-seq data. All methods are evaluated on one semi-simulated, one simulated, and two real datasets resembling several experimental scenarios: globally visible differences in various magnitudes, patient-specific effects on paired data, highly zero-inflated marker expressions, and an immune dataset composed of multiple cell types (see Figure 1C).

In addition, we present CYANUS (CYtometry ANalysis Using Shiny), a user-friendly R Shiny App available at https://exbio.wzw.tum.de/cyanus/. In contrast to existing cytometry analysis platforms like Cytobank (***Kotecha et al.*** (***2010***)) or OMIQ (***Belkina et al.*** (***2019***)), we provide a free platform allowing researchers to analyze normalized, gated cytometry data. To this end, we integrated state-of-the-art methods from CATALYST (***Crowell et al.*** (***2021***)) for preprocessing, visualization, and clustering. Additionally, we integrated those methods for differential marker expression and abundance which showed good performance in our benchmark.

## Results

In this study, we compared existing and novel approaches for detecting differentially expressed markers in CyTOF datasets. The diffcyt package (***Weber et al.*** (***2019***)) employs LMM and limma, which both use median marker expressions per sample and cluster when comparing conditions. In contrast, the methods from the CytoGLMM package (***Seiler et al.*** (***2021***)) make use of the whole data by modeling the condition with all marker expression values (per sample and cluster).

We hypothesize that when reducing the datasets to their medians as in diffcyt, simple statistical tests such as the Wilcoxon rank-sum/signed-rank test, Kruskal-Wallis or (paired) t-test could be effective. We further use a univariate logistic regression to examine whether the CytoGLMM approach could be simplified. To explore whether using the entire distribution of the dataset is beneficial, we modeled the expression data by fitting a zero-inflated beta distribution (BEZI) and a zero-adjusted gamma distribution (ZAGA), respectively. We further used the Earth Mover’s Distance to compare normalized distributions for each marker (and cluster) between groups (CyEMD). For more details, please refer to the Methods section.

We used four different datasets to evaluate method performance for different data distributions. The semi-simulated data contains a clear, globally visible artificial signal for four markers. In the simulated CytoGLMM dataset, five markers are differentially expressed but the differences are only present on patient-level, not overall. The dual platelet dataset contains strong zero-inflation for two platelet activation markers, leading to a median marker expression of zero. The PBMC dataset contains different cell types which is why a differential expression analysis should only be done cell type (cluster)-wise. The method performance will be discussed for each dataset to show strengths and weaknesses of the algorithms. Statistical test may report significant differences that are not meaningful due to their negligible effect size. To account for this, we computed, for all results, the overall (global) and grouped (accounting for patients or other groups) effect size. It should be noted that the grouped effect size must be treated with caution due to the small number of samples (see Methods).

Figure 1 shows a schematic representation of a differential analysis workflow for cytometry data as well as an overview of the methods and datasets investigated in this study.

### Semi-Simulated COVID-19 Dataset With Clean, Globally Visible Difference Between Conditions

For the semi-simulated COVID-19 platelet dataset, we expect to find the markers CD63, CD62P, CD107a, and CD154 differentially expressed between stimulated and non-stimulated samples because an artificial signal was only created for these markers specifically. In the original experiment, platelet expression from baseline samples was compared to expression measured for activated platelets after stimulation with thrombin receptor-activating peptide (TRAP). The four markers whose expression was used for creating the signal, hereinafter referred to as state markers, are known platelet activation markers (***Blair et al.*** (***2018a***)).

All other markers (i.e. type markers) detected by any method can be classified as false positives, since the baseline expression values were not modified.

To examine the sensitivity of the methods, we reduced the differences in expression between the baseline and the spike condition step-wise via a parameter *α* (Equation 1) which indicates by what percentage the difference between the spiked-in expression and the baseline expression is reduced. In the datasets where *α* was set to 1, no marker should be classified as differentially expressed because the spiked-in expressions are equal to their corresponding baseline cell.

The differences are visible on a global level, as we can observe from the overall effect size which is large for CD63, CD62P, CD107a, and moderate for CD154. While all of the other 18 markers have a negligible overall effect size, six of them have a small grouped effect size and one has a moderate grouped effect size. Supplemental Figures 1 and 2 show the results containing all downsampled datasets for activation (state) markers and other (type) markers, respectively.

Table 1 gives an overview of the methods’ performance across all COVID-19 datasets measured by the F1-score. Sensitivity, specificity, and precision on the same datasets can be found in Supplemental Tables 1, 2, and 3, respectively. The methods relying on the median marker expression tend to perform better with an increasing number of cells. The opposite is the case for both methods from the CytoGLMM package, as well as BEZI, ZAGA, and the logistic regression.

**Table 1.** Methods’ performance measured by F1 scores on the semi-simulated COVID-19 dataset. The means and standard deviations of the scores are reported across the multiple *α* values.

| Number of Cells | 1,000 | 2,000 | 5,000 | 10,000 | 15,000 | 20,000 | 4,052,622 |
| --- | --- | --- | --- | --- | --- | --- | --- |
| diffcyt-DS-limma | 0.89 +/- 0 | 0.89 +/- 0 | **1 +/- 0** | **1 +/- 0** | **1 +/- 0** | **1 +/- 0** | **1 +/- 0** |
| diffcyt-DS-LMM | 0.62 +/- 0 | 0.67 +/- 0 | 0.73 +/- 0 | 0.73 +/- 0 | 0.73 +/- 0 | 0.73 +/- 0 | 0.73 +/- 0 |
| t-test | 0.89 +/- 0 | 0.89 +/- 0 | **1 +/- 0** | **1 +/- 0** | **1 +/- 0** | **1 +/- 0** | **1 +/- 0** |
| Wilcoxon test | 0.96 +/- 0.07 | 0.85 +/- 0.07 | 0.96 +/- 0.07 | 0.96 +/- 0.07 | 0.96 +/- 0.07 | 0.96 +/- 0.07 | 0.96 +/- 0.07 |
| Kruskal-Wallis test | **1 +/- 0** | **1 +/- 0** | **1 +/- 0** | **1 +/- 0** | **1 +/- 0** | **1 +/- 0** | **1 +/- 0** |
| CytoGLM | 0.79 +/- 0.05 | 0.69 +/- 0.14 | 0.5 +/- 0.07 | 0.47 +/- 0.09 | 0.48 +/- 0.13 | 0.48 +/- 0.12 | 0.4 +/- 0.05 |
| CytoGLMM | 0.74 +/- 0.06 | 0.47 +/- 0.03 | 0.38 +/- 0.02 | 0.36 +/- 0.03 | 0.34 +/- 0.04 | 0.33 +/- 0.03 | 0.37 +/- 0.04 |
| logRegression | **1 +/- 0** | **1 +/- 0** | **1 +/- 0** | **1 +/- 0** | **1 +/- 0** | **1 +/- 0** | 0.89 +/- 0 |
| ZAGA | 0.96 +/- 0.07 | 0.85 +/- 0.07 | 0.96 +/- 0.07 | 0.93 +/- 0.08 | 0.96 +/- 0.07 | 0.85 +/- 0.07 | 0.89 +/- 0 |
| BEZI | 0.93 +/- 0.08 | 0.96 +/- 0.07 | 0.96 +/- 0.07 | 0.88 +/- 0.16 | 0.96 +/- 0.07 | 0.58 +/- 0.06 | 0.28 +/- 0.09 |
| CyEMD | **1 +/- 0** | **1 +/- 0** | **1 +/- 0** | **1 +/- 0** | **1 +/- 0** | **1 +/- 0** | **1 +/- 0** |

The diffcyt methods can find all activation markers regardless of sample size and signal intensity.

In the negative controls, both methods find markers in the small downsampled datasets (1000 and 2000 cells per patient).

Regarding the statistical tests on expression medians, the Kruskal-Wallis test correctly detects all of the state markers and none of the type markers across all sample sizes and *α* values. The Wilcoxon signed-rank test misses CD154 for *α*=0 regardless of the sample size. In the negative controls, the Wilcoxon test and the t-test find one type marker for n=2000 and the t-test finds one type marker for n=1000. This observation can be made for all *α* values, except for *α*=1.

The CytoGLMM methods find many false positive type markers across all *α* values (except for *α*=1). The number of false positives rises with increasing sample size. For the downsampled datasets, more type markers are found for higher *α* values. Additionally, CD154 cannot be detected by both CytoGLMM methods for *α*=0, as well as by CytoGLM for *α*=0.25.

BEZI fails to find different subsets of the state markers across all sample sizes and *α* values, either due to convergence errors (for datasets bigger than 5000 cells/patient) or because they did not pass the significance threshold of 0.05. Additionally, BEZI classifies PEAR as differentially expressed for all *α* values in the datasets that were not subsampled. ZAGA and the univariate logistic regression also find PEAR in this dataset but not for *α*=1. Similar to the CytoGLMM methods, ZAGA fails to find CD154 for smaller datasets when *α* is set to 0.25. Apart from that, ZAGA and the univariate logistic regression do not make any further false predictions.

Finally, CyEMD was able to classify all markers correctly.

### Simulated Data from CytoGLMM Package With Differences Only Visible on Patient-Level

The design of the CytoGLMM simulation leads to patient-wise differences that are not visible globally. The data was simulated in such a way that m01-m05 are differentially expressed and therefore expected to be found. We observe that indeed, the grouped effect size is large for markers m01-m05 while the overall effect size is negligible (see Supplemental Figure 3).

Table 2 shows an overview of performance measurements on all subsets of this dataset. For more detailed results, we refer to Supplemental Figure 3.

**Table 2.** Methods’ performance on the simulated CytoGLMM dataset. Sensitivity, specificity, precision, and F1 score are shown for each method. Means and standard deviations of the scores are reported across the multiple numbers of cells. If no positive classification was made, precision and F1 score cannot be computed and are marked as NaN in the table.

|  | Sensitivity | Specificity | Precision | F1 Score |
| --- | --- | --- | --- | --- |
| diffcyt-DS-limma | 0.97 +/- 0.08 | 0.99 +/- 0.03 | 0.98 +/- 0.06 | 0.97 +/- 0.05 |
| diffcyt-DS-LMM | **1 +/- 0** | 0.96 +/- 0.04 | 0.9 +/- 0.09 | 0.95 +/- 0.05 |
| t-test | 0.97 +/- 0.08 | **1 +/- 0** | **1 +/- 0** | 0.98 +/- 0.04 |
| Wilcoxon test | 0.49 +/- 0.34 | **1 +/- 0** | **1 +/- 0** | 0.81 +/- 0.08 |
| Kruskal-Wallis test | 0 +/- 0 | **1 +/- 0** | NaN | NaN |
| CytoGLM | 0.8 +/- 0.38 | **1 +/- 0** | **1 +/- 0** | 0.96 +/- 0.1 |
| CytoGLMM | 0.97 +/- 0.08 | 0.97 +/- 0.05 | 0.94 +/- 0.12 | 0.95 +/- 0.07 |
| logRegression | **1 +/- 0** | **1 +/- 0** | **1 +/- 0** | **1 +/- 0** |
| ZAGA | 0.91 +/- 0.16 | 0.91 +/- 0.12 | 0.83 +/- 0.19 | 0.85 +/- 0.14 |
| BEZI | 0.86 +/- 0.15 | 0.93 +/- 0.07 | 0.84 +/- 0.16 | 0.83 +/- 0.1 |
| CyEMD | 0 +/- 0 | **1 +/- 0** | NaN | NaN |

The two methods that cannot perform a paired analysis, CyEMD and the Kruskal-Wallis test on marker expression medians, do not find any marker to be differentially expressed. Consequently, they achieve a specificity of 1 and all other measurements are 0 or undefined. Overall, the diffcyt methods have a high performance and gain power for greater numbers of cells. This effect can also be observed for most other methods. CytoGLMM’s and CytoGLM’s scores are close to 1 except for the sensitivity scores for CytoGLM which vary more strongly since some of the differentially expressed markers cannot be detected for low cell counts. BEZI and ZAGA lose performance mostly because the algorithms do not converge. Apart from that, they yield high scores. Only the univariate logistic regression can correctly identify all differentially expressed markers without a false positive discovery.

### Dual Platelet Dataset With Zero Median Marker Expression

This dataset was generated by collecting two samples from each participant and stimulating one of the two samples with TRAP to activate the platelets. Therefore, we expected to find platelet activation (state) markers like CD63, CD62P, CD154 and CD107a to be differentially expressed between the two conditions. Figure 4C shows that CD154 and CD107a have a median marker expression of zero, posing a challenge for the methods using only marker medians.

We tested our methods twice on this dataset. For the first run, the patient ID was included as a grouping variable while the second analysis was unpaired (see Figure 2). We used the Wilcoxon rank-sum test and the Wilcoxon signed-rank test in the unpaired and paired design, respectively.

**Figure 2.**
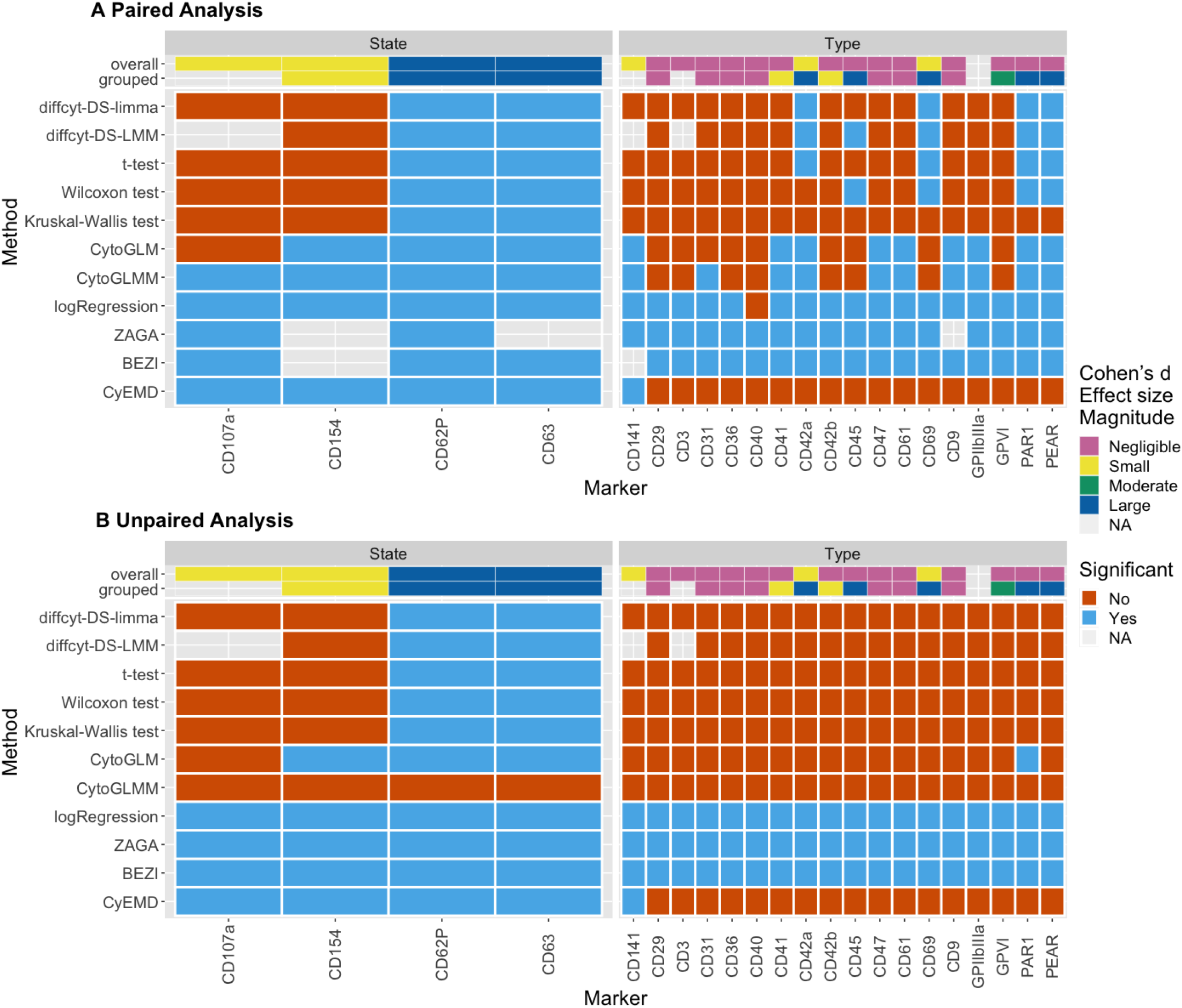
Method results for the dual dataset with patient id as grouping variable **(A)** and without any grouping variable **(B)**. Results colored in blue if the adjusted p-value < 0.05, else in red. Uncolored tiles mean convergence errors of the method for the specific marker. The overall and grouped effect size magnitudes per marker are shown at the top. Overall effect size refers to *Cohen’s d* magnitudes using all expression data between two conditions. The magnitudes indicated by grouped effect size are computed in a paired fashion on the median marker expressions per sample. Wilcoxon test refers to the Wilcoxon signed-rank test and the Wilcoxon rank-sum test for the paired and the unpaired analysis, respectively. Markers are divided into their marker class (state and type).

ZAGA, BEZI, and the univariate logistic regression classify all markers as significant or do not converge. The issues of these three methods are examined thoroughly in the discussion.

The five algorithms (diffcyt-limma, diffcyt-LMM, t-test, Wilcoxon test, and Kruskal-Wallis test) using only median expressions find the two state markers CD63 and CD62P but are not able to find the zero-inflated markers CD154 and CD107a. In the unpaired run, no type markers are found by these methods. In the analysis with patient ID as grouping variable, the Wilcoxon signed-rank test, t-test, and both diffcyt methods find PAR1, PEAR, and CD69 to be significantly differentially expressed between the non-stimulated and stimulated samples. CD42a was also found by the t-test and both diffcyt methods. Each of the four markers has a large grouped effect size.

Two of the three methods using whole marker expression, CyEMD and CytoGLMM, classify all four state markers as significant. CytoGLM only misses CD107a which has a small overall effect size. Since CytoGLMM cannot be run without a random effect, its result for this run is not reliable. Looking at the results for the type markers, CyEMD finds CD141 and CytoGLM finds PAR1 when no paired analysis is performed. After including the patient ID as grouping variable, additional markers are found by all methods able to incorporate this information. CytoGLMM, CytoGLM, and CyEMD all detect CD141 (small overall effect size). The two methods of the CytoGLMM package find several additional markers: CD41, CD61, PAR1, GPIIbIIIa, CD141, CD9, PEAR, CD47, CD31, and CD42a. While PAR1, PEAR, and CD42a are also found by other methods (as mentioned above), some of these markers (CD61, CD47, CD9, and CD31) have negligible effect sizes which is why we classify them as false positives (see Supplemental Figure 4).

### PBMC Dataset With Different Cell Types

Since our first real dataset, the dual platelets dataset, only contains one cell type, we also evaluated the different approaches on the PBMC dataset by ***Bodenmiller et al.*** (***2012***) which contains eight immune cell types annotated by ***Nowicka et al.*** (***2019***). For each cluster of cell types, differential expression analysis was performed, comparing the reference condition against the cells that were cross-linked with B cell receptor/Fc receptor (BCR/FcR-XL). We expected to find pS6 differentially expressed since these findings have been made in the original paper (***Bodenmiller et al.*** (***2012***)). Supplemental Figure 5 shows an overview of the results.

Many markers were significant across all clusters. Independent of the method, overall and grouped effect sizes were large for numerous markers in all clusters (see Supplemental Figure 5).

Of all possible 192 marker-cluster combinations (24 markers in 8 cell types), the univariate logistic regression, BEZI, and ZAGA find the most markers to be differentially expressed (168, 156, and 155, respectively).

The methods that are not able to include a grouping variable, CyEMD and the Kruskal-Wallis test, find the least markers to be differentially expressed (86 and 88, respectively). In contrast to the dual dataset results, CytoGLMM and CytoGLM do not produce more positive predictions than the statistical tests, CyEMD, or the diffcyt methods.

### Runtime

The runtimes for the complete datasets are shown in Table 3. For the runtimes of the subsampled datasets, please refer to Supplemental Figure 6.

**Table 3.**
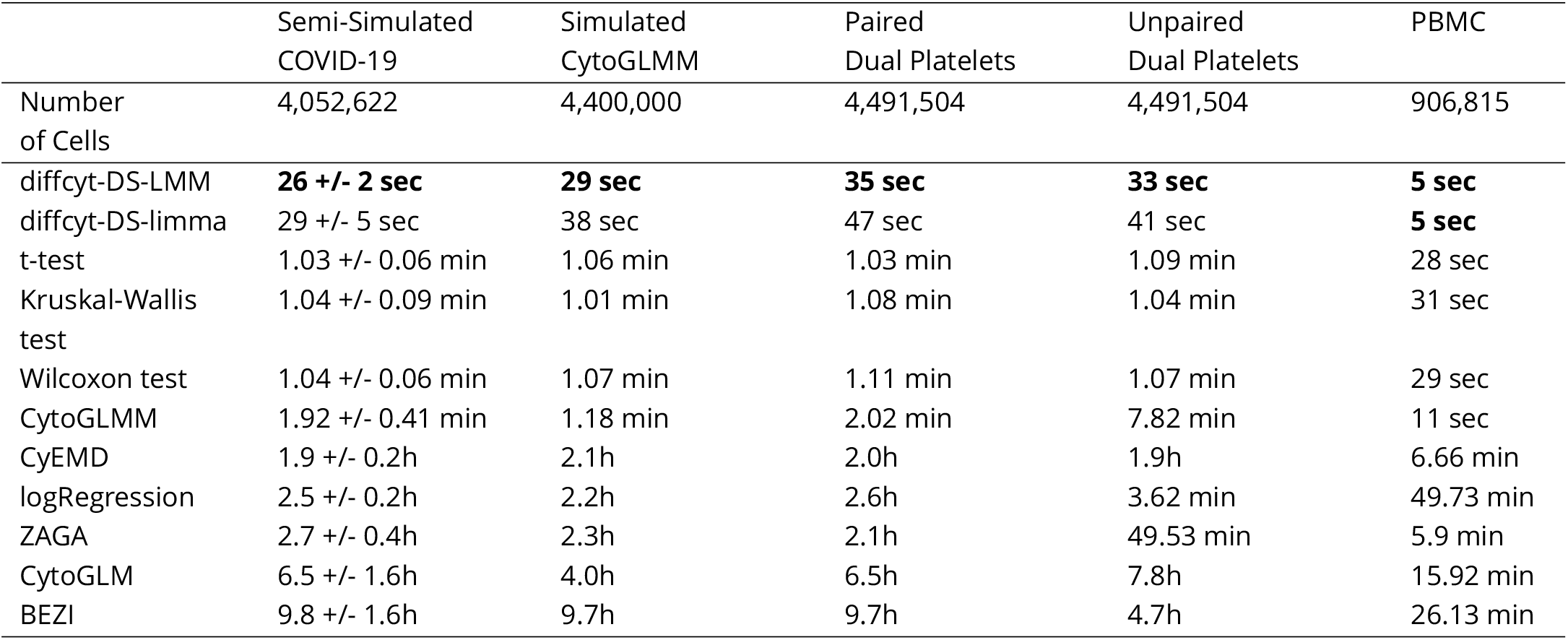
Runtime of the methods on the different datasets. The methods that reduce the data to medians and CytoGLMM have very low runtime requirements while CytoGLM and BEZI are slow on big datasets. The univariate logistic regression, CyEMD, and ZAGA have moderate runtimes.

The diffcyt methods outperform all other methods in terms of runtime. All in all, methods that use median marker expressions are fast independent of sample size. CytoGLMM and the unpaired logistic regression are quick as well, even though they take the whole distribution into account.

The paired univariate logistic regression, CyEMD, and ZAGA have moderate runtimes while CytoGLM and BEZI often run more than six hours on the big datasets.

## Discussion

### Semi-Simulated COVID-19 Dataset With Clean, Globally Visible Difference Between Conditions

Regarding the grouped effect sizes, we can observe a problem that occurs when the differences between the groups are very small, yielding a standard deviation close to zero (see Equation 3). By dividing by a value close to zero, bigger values for the effect sizes are obtained, hence seven of the markers that were not spiked in appear to have a small or even moderate grouped effect size. The difference in means between the two conditions is not significant, as shown by the paired t-test. Therefore, we recommend checking the effect size when a marker is classified as differentially expressed and additionally, checking the results of the paired t-test when the grouped effect size is not negligible because both methods compare paired means.

The diffcyt methods and the statistical tests perform well, especially for higher sample sizes. We hypothesize that the markers that are found in the small negative control datasets were detected because of noise in the measured data. Due to the law of large numbers, the median becomes more reliable for higher cell counts. Therefore, methods that reduce the expression data to medians become more stable with growing dataset size.

The CytoGLMM methods produce a high number of false positives for all *α* values except for *α*=1, especially with rising sample size. A possible explanation could be that the multivariate generalized mixed effect models become too sensitive to small changes when there are only few bigger differences (here, CD62P and CD63) because all markers are included as explanatory variables. Therefore, the condition is modeled as a result of various small changes which are present because of the semi-simulated nature of the data. The increasing sample size seems to reduce the magnitude of the p-values.

The Wilcoxon signed-rank test and the CytoGLMM methods miss CD154 for *α*=0 and *α*=0.25 but find it for the other *α* values because of the way the artificial signal was created. Looking at the medians per patient for the full number of cells (see Supplemental Figure 7), it is visible that the medians of the spike condition are higher for *α*=0.25, 0.5, and 0.75 than for *α*=0 in two patients. Additionally, the medians are extremely close to zero. Due to this strong zero-inflation, the spiked-in expressions for CD154 were sometimes smaller than the baseline expressions. As Equation 1 leads to a convergence of the activated measurements towards the baseline measurements, the spiked-in values become higher for the cells where the activated measurements were smaller than the baseline measurements. Therefore, the expression is less zero-inflated in these two patients and the marker can be found by the three methods for higher *α* values.

For BEZI, we see its high sensitivity for big sample sizes (see dual dataset), since PEAR is found across all *α* settings in the big dataset, while it is not found for the downsampled datasets. ZAGA and the univariate logistic regression yield reliable results for datasets with a clean, globally visible difference, especially for smaller sample sizes.

### Simulated Data from CytoGLMM Package With Differences Only Visible on Patient-Level

Because this data is paired, we expect that only methods that can handle paired data can detect the differentially expressed markers between the two conditions. This is confirmed as all methods except for CyEMD and the unpaired Kruskal-Wallis test detect the differential expression and can be sensitive to small changes in expression that are only detectable at the patient level.

CytoGLMM’s false detection of one marker suggests an over-sensitivity further described in the next section.

### Dual Platelet Dataset With Zero Median Marker Expression

The results for this dataset clearly show the problem of reducing the data on median marker expressions to perform differential expression analysis. Methods taking the whole marker expression into account find markers with zero-median marker expression, whereas methods working on the medians are not able to find these.

Furthermore, the results differ depending on the applied method. The markers PAR1 and PEAR are found by several methods. While PEAR has a higher expression in the stimulated condition, PAR1 is less expressed in this condition (see Supplemental Figure 8). In literature, the PEAR receptor has been described to be increased on the platelet membrane after stimulation with several activators (***Kauskot et al.*** (***2012***)), while the effect on PAR1 expression after stimulation depends on the agonist. Studies using a PAR1-AP are in line with our findings and show a decreased amount of the PAR1 receptor on the platelet surface after stimulation (***Ramström et al.*** (***2008***)).

CD69 which is found by limma, LMM, the Wilcoxon test, and the t-test, shows a higher signal after stimulation. Several studies have observed a similar trend for CD69 increase upon stimulation (***Testi et al.*** (***1990***, ***1992***)). CD42a is detected by the two diffcyt methods, the CytoGLM/M methods, and the t-test, and shows a decreasing trend after TRAP stimulation. This also has been previously shown in platelets using CyTOF (***Blair et al.*** (***2018b***)). Several other studies examined a decrease of CD42a expression after stimulation with activators ADP (***Braune et al.*** (***2014***)) and collagen (***Hagberg et al.*** (***1997***)). The biological reason behind the differential expression of the two markers CD141 and CD45 remains unclear. In general, CD141 is not found to be expressed on platelets (***Bongiovanni et al.*** (***2021***)) whereas CD45 has shown to be present on the surface of several platelets (***Gabbasov et al.*** (***2014***)).

The application of ZAGA, BEZI, and the univariate logistic regression is unfeasible for a real dataset of this size. CytoGLMM and CytoGLM produce at least three false positives due to their high sensitivity. Additionally, CytoGLM misses one of the two highly zero-inflated activation markers. The diffcyt methods, the t-test, and the Wilcoxon signed-rank test perform fast and yield reliable results but miss the two activation markers that have a median of zero. The Kruskal-Wallis test performs worse than the Wilcoxon signed-rank test on this dataset because it is not able to handle paired data and could therefore not detect markers like PAR1, PEAR, CD69, or CD42a. Lastly, CyEMD detects the globally visible changes for the activation markers and CD141 but fails to detect any of the changes that can only be seen on the patient level as seen in Figure 2.

### PBMC Dataset With Different Cell Types

The evaluation of this dataset is limited by the number of cells per sample and cluster (see Supplemental Figure 9). For cell types with less than 1000 cells per sample, noise is distorting the analysis.

When ***Weber et al.*** (***2019***) evaluated their diffcyt methods on this dataset, they could confirm that pS6 is differentially expressed in B-cells. From Figure 5, it becomes apparent that all methods can find this marker. Moreover, ***Nowicka et al.*** (***2019***) showed that the diffcyt methods identify pS6 to be also differentially expressed in other cell types. All methods tested in this study confirm this finding and find pS6 differentially expressed in all cell types except for dendritic cells.

In contrast to the dual platelet dataset, the univariate logistic regression, BEZI, and ZAGA were not suffering from a clear over-identification of markers. Because the PBMC dataset is rather small (172,791 cells in total vs. 4,491,504 in the dual platelet dataset), we hypothesize that the higher the number of cells, the less suitable these three methods become. This is due to the influence of large sample sizes on the magnitude of the p-values (***Lin et al.*** (***2013***)).

Compared to the dual platelet dataset, the CytoGLM/M methods did not identify more markers as significantly differentially expressed than the other methods, even though there are more markers with a large effect size.

### Conclusion and Outlook

Existing approaches for differential marker expression analysis were compared with simple and advanced novel approaches that rely either on median or on full marker expression data using two real, one semi-simulated, and one simulated dataset.

A limitation on the level of dataset evaluation is that we could not interpret the results obtained on the PBMC dataset biologically. We could therefore not describe which markers were falsely classified as differentially expressed and which markers were overlooked. Additionally, we did not include a dataset with batch effects but assumed that the data had already been corrected for it. Theoretically, it should be possible to include a batch effect as a random effect or additional term in a model. This can be done for all the approaches we evaluated but the statistical tests and CyEMD. Finally, the downsampling of the spiked and the CytoGLMM datasets was not repeated multiple times. If that would have been done, the results would be more reliable and robust. In this study, repeating the evaluations that many times was not feasible because of the high runtime requirement of BEZI and ZAGA.

All in all, the diffcyt methods perform fast and yield good, trustworthy results when the median of the differentially expressed marker is unequal to zero. Nevertheless, they did not outperform a simple, Wilcoxon signed-rank test or t-test on the medians, meaning that a more complicated model is not certainly necessary to detect significant differences in CyTOF marker medians. The comparison with the Kruskal-Wallis test on marker medians shows that the clear advantage of the Wilcoxon/t-test is the ability to compute a paired test statistic.

Regarding the cytoGLMM methods, we observe that small, individual changes can be detected as well as globally visible changes on very clean data, even when it is strongly zero-infated. Additionally, cytoGLMM is fast even though it takes the whole distribution into account. On the other hand, the methods are extremely sensitive to changes even without a small, grouped effect size and classify many markers to be differentially expressed, especially with growing dataset size. Therefore, we recommend checking for overlaps between cytoGLMM and other methods, making diagnostic plots and looking at the effect size magnitude when running cytoGLMM on larger, real datasets.

BEZI, ZAGA, and the univariate logistic regression proved to be infeasible for larger, real datasets. While the performance on the completely artificial CytoGLMM dataset was acceptable, the method performance dropped for the semi-simulated spike dataset and eventually only produced positive predictions on the real, dual platelets dataset. Additionally, BEZI is unacceptably slow.

Finally, our novel method CyEMD exploits the advantages of taking the whole marker expressions into account and still performs well on big datasets because it partitions the distribution into bins and computes p-values via permutation tests. We showed that the EMD approach can detect differentially expressed markers that are strongly zero-inflated in an acceptable amount of time. Additionally, the approach should be able to find differences in bimodal or skewed marker expressions, even when the medians are similar. A disadvantage to the EMD approach is that it cannot detect differentially expressed markers when the changes are only visible by comparing expressions group- or patient-wise.

Our results across datasets with different properties show that each of the tested methods comes with its own strengths and weaknesses. Taking factors like runtime, zero-inflation and skewness and sample groups into account, we offer a guideline for users to choose optimal methods for their analysis (Figure 3). However, often several methods are suitable for a given scenario and should be compared to obtain robust and interpretable results.

**Figure 3.**
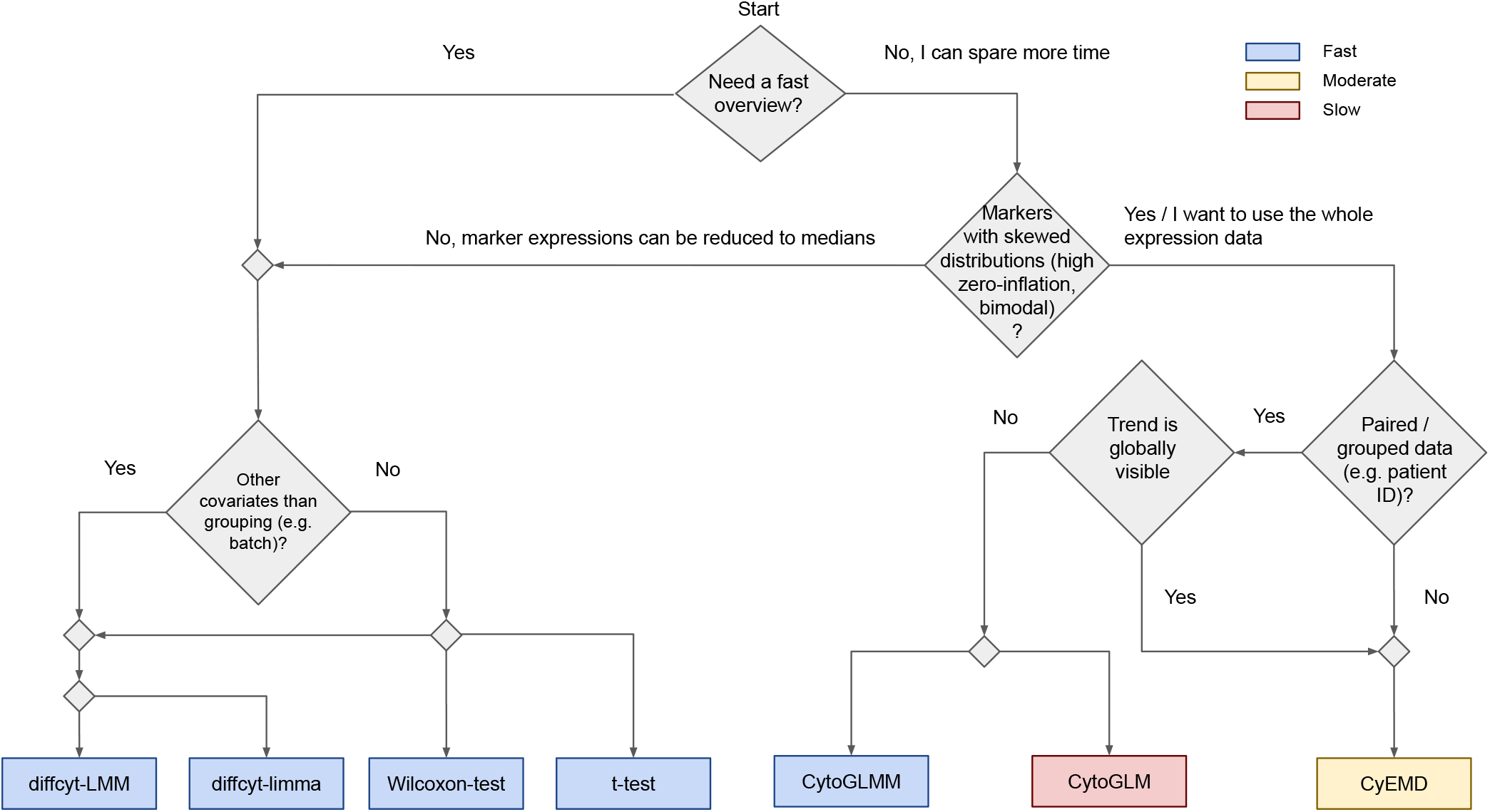
Overview of the methods suitable for CyTOF data. Several scenarios can occur while analyzing CyTOF data. This graph helps to identify the most suitable method and includes the runtime of the different methods.

To make such a comparative analysis easily accessible, we integrated the diffcyt methods, the Wilcoxon rank-sum and signed-rank test, the t-test, the cytoGLMM methods, and CyEMD into a user-friendly R Shiny App CYANUS available at https://exbio.wzw.tum.de/cyanus/. CYANUS (CYtometry ANalysis Using Shiny) allows the user to analyze gated and normalized cytometry data (i.e. flow cytometry as well as CyTOF) with state-of-the-art methods from CATALYST (***Crowell et al.*** (***2021***)). For differential abundance analysis, we integrated the methods included in the diffcyt package. All differential analysis methods can be easily compared to each other, enabling thorough analysis of cytometry data exploiting the advantages of the various approaches.

## Methods

### Data Description

For the evaluation of the differential expression methods, we worked with four different datasets. The methods were tested on one semi-simulated, one simulated, and two real CyTOF datasets (see Figure 1).

#### Semi-Simulated COVID-19 Data

The semi-simulated COVID-19 dataset originates from the University Hospital rechts der Isar, Munich, Germany (***Bongiovanni et al.*** (***2021***)). The original dataset comprises CyTOF data of 8 symptomatic SARS-CoV-2-infected patients, hospitalized between March and May 2020. Additionally, 11 healthy donors were included in the study. A baseline sample (non-stimulated platelets) and one sample stimulated with TRAP was prepared for each donor.

In order to study the sensitivity of the methods to changes in the expression patterns, we performed the following data generation procedure. Firstly, the baseline healthy samples were randomly split in half. Half of a sample was used for randomly spiking in the expression values for the four known activation markers (CD62P, CD63, CD107a, CD154) from the activated sample of the corresponding patient. Because this leads to very clear, well distinguishable results, we reduced the differences in expression between baseline and spike expressions for the four markers using the following formula:

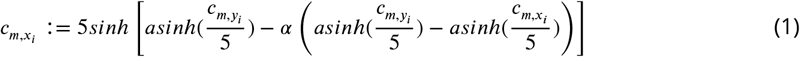

where *m* is the marker, *c_m,x_i__* is the raw value measured for the baseline sample for cell *x_i_, c_m,y_j__* is the raw value measured for the activated sample for cell *y_j_*, *X* = *x*_1_, …, *x*_*N*/2_ are the indices of the baseline cells whose expression was randomly replaced and *Y* = *y*_1_, …,*y*_*N*/2_ are the indices of the activated cells whose values were used for spiking. Since we wanted to observe the differences in the asinh transformed expression values, the reduction was made on the level of the transformed values. Using the formula, five datasets were produced by setting *α* to 0 (full intensity), 0.25, 0.5, 0.75, and 1.0 (control) (see Figure 4B). Each dataset contains eleven paired samples with 4,052,622 cells in total (see Supplemental Table 4 for the number of cells per sample). This approach was inspired by the diffcyt benchmarking strategy (***Weber et al.*** (***2019***)). In contrast to their approach, we did not use differences in means and standard deviations between the two conditions for reducing the signal but the actual differences between *c_m,x_i__* and *c_m,y_j__*.

**Figure 4.**
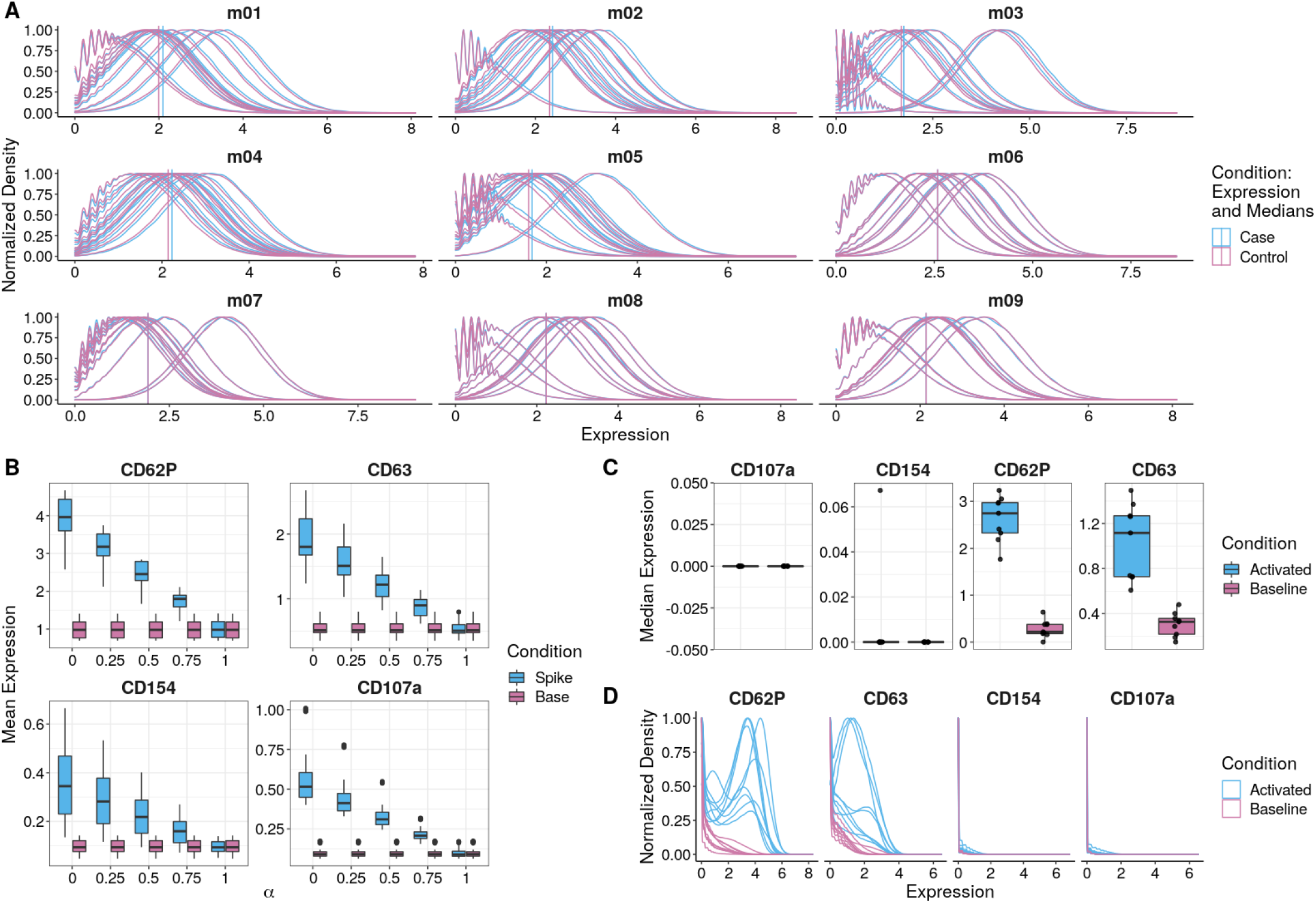
Marker Expressions of the simulated CytoGLMM (A), semi-simulated COVID-19 (B), and the dual platelet datasets (C,D). **(A)** Normalized density of the markers m01-m09 of the dataset simulated using the CytoGLMM data generation process by ***Seiler et al.*** (***2021***). The markers m01-m05 are simulated to be differentially expressed in such a way that the expression differs slightly but consistently for each patient. Meanwhile, the median marker expressions of the whole dataset, marked by the vertical lines, do not differ significantly. **(B)** Mean expressions for the four spiked-in activation markers at different intensities. For *α*=0 (full intensity), the originally measured expressions of the corresponding activated sample were used. Subsequently, *α* was repeatedly increased by 0.25 in order to reduce the difference between the spiked and the base condition so that the differences would become harder to detect. *α*=1 was used as control dataset. **(C)** Median expression of state markers of the dual platelet dataset. Markers CD62P and CD63 are higher expressed in the activated condition. The median marker expression of CD107a and CD154 is zero, except for one sample. **(D)** Normalized density of state markers of the dual platelet dataset. CD107a and CD154 show a small difference in the expression.

#### Simulated CytoGLMM Data

To investigate the methods’ handling of data without global but paired differences in expression, we used a customized version of the data simulation process described by ***Seiler et al*.** (***2021***). The algorithm samples from a Poisson GLM with an underlying hierarchical model combining effects on cell and donor-level for two conditions. Figure 4A shows that this leads to expression differences on a patient level, but not overall. The resulting simulated dataset used in this study consists of 20 markers, of which 5 are differentially expressed, in 22 paired samples from 11 patients with 200,000 cells per sample.

#### Dual Platelet Data

We used a CyTOF platelet dataset originating from the University Hospital rechts der Isar, Munich, Germany, consisting of platelet heterogeneity measurements of patients with chronic coronary syndrome receiving dual anti-thrombotic therapy. The dataset contains 4,491,504 cells and includes 18 paired samples from 9 donors in two conditions: non-stimulated and stimulated (TRAP). For the exact number of cells per sample, refer to Supplemental Table 5. The panel containing 22 protein markers (see Supplemental Table 6) includes four well-known platelet activation markers (***Blair et al***. (***2018a***)). Two of the platelet activation markers, CD63 and CD62P, are known to be highly upregulated after TRAP stimulation, whereas CD107a and CD154 are upregulated less strongly (see Figures 4C, D).

#### PBMC Data

The peripheral blood mononuclear cells (PBMCs) dataset originating from ***Bodenmiller et al*.** (***2012***) consists of samples from 8 healthy donors in 12 conditions. ***Nowicka et al.*** (***2019***) performed a complete CyTOF analysis on a subset of this data containing the reference and one stimulated condition. In the stimulated condition, the cells were cross-linked with B cell receptor/Fc receptor for 30 minutes. This subset consists of 172,791 cells in 16 paired samples from 8 patients (see Supplemental Table 7). ***Nowicka et al.*** (***2019***) manually merged 20 clusters obtained via meta clustering into 8 cell populations which were made publicly available. In this study, this annotated and well-described subset was used.

#### Downsampling of Artificial Datasets

To review changes in the methods’ power and runtime with respect to sample size, we downsampled the two simulated datasets. Both the spiked COVID-19 and the simulated CytoGLMM data was subsampled to contain 1000, 2000, 5000, 10000, 15000, and 20000 cells per patient. For both datasets, we sampled in such a way that the smaller sets are always subsets of the bigger ones. The same cells were used in the COVID-19 dataset for different *α* values, to ensure a fair comparison.

#### Effect Size

To quantify the difference between marker expressions, we computed *Cohen’s d* (***Cohen*** (***1977***)) for each marker in every dataset using the `rstatix` R package (***Kassambara*** (***2021***)). The thresholds for the absolute value of *d* to consider the magnitude of the effect size at least *small, moderate,* and *large* are 0.2, 0.5, and 0.8, respectively. Values smaller than 0.2 are referred to as *negligible.* The effect size was calculated overall (on the whole expression) and grouped (based on the median marker expression of the paired samples). The overall effect size compares marker intensities between two conditions by using their mean and (shared) standard deviation:

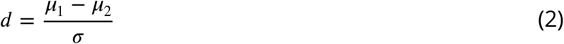

Differences on the patient-level can be captured with a paired effect-size estimation, defined as grouped effect size. To obtain paired data points, the expression median was computed for each sample. Additionally, the paired effect-size allows us to check whether significant results from the paired t-test have considerable effect-sizes and whether effect-sizes of higher magnitudes are statistically significant because both methods investigate differences in population means. Since the sample size for the paired calculation is limited to the number of patients *n,* which is smaller than 20 for all datasets, and there is a known upwards bias for small sample sizes, we used Hedges’ correction to adjust for that. The grouped effect size with Hedges correction is computed as follows (***Hedges and Olkin*** (***1985***)):

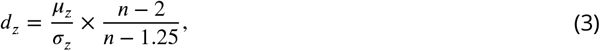

where *x* and *y* are the median marker expressions of two groups with paired samples and *z* is their difference *z* = *x* – *y*.

### Differential Analysis

In this work, we compared the existing approaches for differential marker expression analysis from ***Weber et al.*** (***2019***) (diffcyt-limma, diffcyt-LMM) and ***Seiler et al.*** (***2021***) (CytoGLM, CytoGLMM) with simple and advanced novel approaches that rely either on median or on full marker expression data. The simple approaches consisted of a t-test, a Wilcoxon test (both paired and unpaired) on median marker expressions, a Kruskal-Wallis test on median marker expressions, and a univariate logistic regression predicting the condition from the whole marker expression profiles. More advanced approaches comprised modeling the marker expression distributions with a zero-infated beta distribution (BEZI) and a zero-adjusted gamma distribution (ZAGA). Furthermore, we developed CyEMD, a method which compares the normalized distributions using the Earth Mover’s Distance (EMD). All p-values mentioned in this study have been adjusted per method and dataset to control the false-discovery rate using the Benjamini-Hochberg procedure at a significance level of 0.05.

#### Diffcyt Methods

The diffcyt limma method fits a linear model for each marker-cluster combination, predicting the sample medians from the corresponding conditions. The LMM method builds a linear mixed-effects model and can therefore handle random effects in contrast to the limma method where a grouping variable can be included only as an additional fixed effect (***Weber et al.*** (***2019***)). The diffcyt methods can easily incorporate other covariates such as batch effects in their model as additional terms.

#### CytoGLMM Methods

Instead of predicting the expression from the conditions, the CytoGLMM methods fit a generalized mixed model predicting the conditions from the whole expression vectors. The package contains two methods, CytoGLMM and CytoGLM. The former can only handle grouped data since it relies on a random effect like patient ID whereas the latter can also handle unpaired data. CytoGLM builds a bootstrapped generalized linear model while CytoGLMM builds a generalized linear mixed model (***Seiler et al*.** (***2021***)). In this study, 500 bootstrap replications were used. These methods can also include additional terms in their model.

#### Logistic Regression

In order to find out whether the CytoGLMM approach based on the whole marker expression could be simplified, we fitted univariate logistic regression models per marker and cluster and extracted the p-value from the regression model. A multivariate approach was omitted since the markers are not statistically independent by design. CytoGLMM partially evades this problem by fitting a hierarchical model containing random slopes and intercepts for the grouping variable (patient ID) which assumes dependent errors.

#### Approaches Modeling the Expression: BEZI, ZAGA

As CyTOF data can be strongly zero-inflated (***Papoutsoglou et al*.** (***2019***)), we fit a zero-inflated beta distribution (BEZI) as well as a zero-adjusted gamma distribution (ZAGA) to our expression data. As a basis, we chose the gamma distribution for modeling non-zero expressions because it was demonstrated that the gamma distribution fits expression data more often than other non-Normal distributions (***de Torrenté et al.*** (***2020***)). A common choice for single cell RNA-seq data is the negative binomial distribution (***He et al.*** (***2021***)) which is not suitable for CyTOF data as it requires discrete values. Therefore, we selected the beta distribution as a conjugate to negative binomial distribution, i.e. it belongs to the same probability distribution family. To model the marker expression distributions of CyTOF data, the condition was used as an explanatory variable and its model coefficient was tested for equality to zero. For this, we used the gamlss and the gamlss.dist packages (***Rigby and Stasinopoulos*** (***2005***); ***Stasinopoulos and Rigby*** (***2021***)). To model changes on a patient level for paired data, random intercepts can be included.

The zero-adjusted gamma distribution, *ZAGA*(*μ, σ, ν*), is a continuous distribution on (0, ∞). The response variable *Y* ~ *ZAGA*(*μ, σ, ν*) ∈ [0, ∞) is modeled using the mixed probability function *f_y_*(*y*|*μ, σ, ν*):

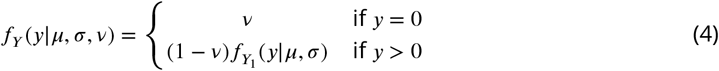

where *y* ≥ 0, *μ* > 0, *σ* > 0, and 0 < *ν* < 1, and where *Y*_1_ ~ *GA*(*μ, σ*). The parameter *ν* is the non-zero probability for *Y* = 0. For *Y* ∈ (0, ∞), *Y* is gamma-distributed.

The zero-inflated beta distribution, *BEZI*(*μ, σ, ν*), is defined on [0, 1). The response variable *Y* ~ *BEZI*(*μ, σ, ν*) ∈ [0, 1) is modeled using the mixed probability function *f_Y_*(*y*|*μ, σ, ν*):

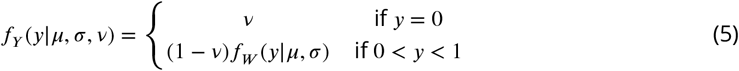

where 0 < *σ* < 1, *μ* > 0, 0 < *ν* < *ν* 1. The beta distribution *f_W_*(*y*|*μ, σ*) is based on the work of ***Ospina and Ferrari*** (***2012***). To fit a zero-inflated beta distribution on CyTOF data, the marker expressions were first scaled to the range [0, 1). For further details regarding the implementation of gamlss, please refer to ***Rigby et al*.** (***2020***).

#### CyEMD

Our novel approach, CyEMD, uses the Earth Mover’s Distance to compare normalized distributions for each marker (and cluster) between groups.

For two normalized histograms *P* and *Q,* the EMD is calculated by minimizing the cost of transforming one into the other. The histograms are represented as *P* = {(*P*_1_, *w*_*p*1_),…, (*p_n_*, *w_pn_*)} and *Q* = (*q*_1_, *w*_*q*1_),…, (*q_n_*, *w_qn_*), where *p_i_*/*q_j_* is the center of the *i*th/*j*th histogram bin and *w_pi_*/*w_qj_* describes the height of the corresponding bin for *P*/*Q*.

To transform histogram P into histogram Q, certain proportions of the bins *p_i_,*, *q_j_* differing between P and Q need to be moved to other bins. The optimization problem for this task is how much has to be transferred from one bin to another bin (defined as f ow *F* = [*f_ij_*]) in order to minimize the cost. The f ow is weighted according to the distances *d_ij_* between the bins such that transporting a high amount of a bin over a long distance is penalized (***Rubner et al*.** (***1998***)):

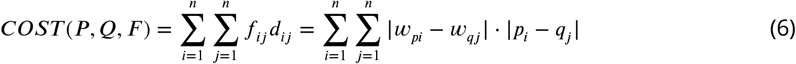

After normalizing the minimal cost by the overall flow we get

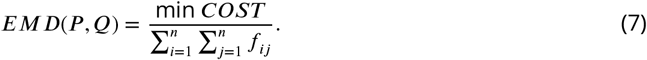

Since the expression densities in CyTOF data can have different ranges for distinct values, we use a f exible bin width estimated by the Freedman-Diaconis rule evaluated on all nonzero values:

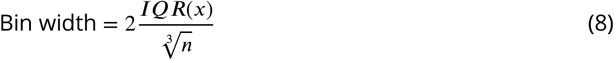

where *IQR*(*x*) is the interquartile range of nonzero marker expressions and *n* is the number of observed expressions (***Freedman and Diaconis*** (***1981***)).

To determine the significance of the EMD between two marker expressions, a permutation test (500 permutations) that permutes the condition labels sample-wise is performed to obtain a p-value for each marker.

As opposed to ***Wang and Nabavi*** (***2018***), we compute the EMD on normalized histograms, which can be done in linear time. In order to speed up the computationally intensive EMD computation, we implemented this part in C++. Furthermore, SigEMD permutes the labels cell-wise instead of sample-wise which proved to be infeasible for big datasets since the empirical p-values become smaller with growing dataset size.

## Data Availability

The scripts for the analysis and the code for the Shiny App are available at https://github.com/biomedbigdata/cyanus under the GPL-3 license.

The original COVID-19 dataset is publicly available at flowrepository.org, accessible at repository ID FR-FCM-Z4AE. The script for producing the semi-simulated COVID-19 data is provided in the Github repository. The simulated CytoGLMM data can be reproduced using a script of the Github repository. Access to the dual dataset (9 patients) is granted upon request. The original PBMC dataset is published at www.cytobank.org/nolanlab. We followed the CyTOF workflow by ***Nowicka et al***. (***2019***) and downloaded the data using HDCytoData (***Weber and Soneson*** (***2019***)). The manual cluster annotation of the CyTOF workflow can be downloaded from http://imlspenticton.uzh.ch/robinson_lab/cytofWorkflow/.

## Acknowledgments

The authors thank Simona Ursu and Sarah Warth at the Core Facility Cytometry of the Ulm University Medical Facility for their support acquiring both platelet datasets. We thank Marc Rosenbaum, Dries van Hemelen, Gloria Martrus and Mayur Bakshi for excellent technical assistance, and Kilian Kirmes for testing and evaluating our app.

