## Supplemental Material for "A Systematic Comparison of Differential Analysis Methods for CyTOF Data"

657 ## Appendix 0

**Appendix 0 Table 1.** Sensitivity of the semi-simulated COVID-19 dataset. The means and standard deviations of the methods are reported across the multiple $\alpha$ values.

| Number of Cells | 1,000 | 2,000 | 5,000 | 10,000 | 15,000 | 20,000 | 4,052,622 |
|---|---|---|---|---|---|---|---|
| diffcyt-DS-limma | **1 +/- 0** | **1 +/- 0** | **1 +/- 0** | **1 +/- 0** | **1 +/- 0** | **1 +/- 0** | **1 +/- 0** |
| diffcyt-DS-LMM | **1 +/- 0** | **1 +/- 0** | **1 +/- 0** | **1 +/- 0** | **1 +/- 0** | **1 +/- 0** | **1 +/- 0** |
| t-test | **1 +/- 0** | **1 +/- 0** | **1 +/- 0** | **1 +/- 0** | **1 +/- 0** | **1 +/- 0** | **1 +/- 0** |
| Wilcoxon test | 0.94 +/- 0.12 | 0.94 +/- 0.12 | 0.94 +/- 0.12 | 0.94 +/- 0.12 | 0.94 +/- 0.12 | 0.94 +/- 0.12 | 0.94 +/- 0.12 |
| Kruskal-Wallis test | **1 +/- 0** | **1 +/- 0** | **1 +/- 0** | **1 +/- 0** | **1 +/- 0** | **1 +/- 0** | **1 +/- 0** |
| CytoGLM | 0.81 +/- 0.12 | 0.88 +/- 0.14 | 0.88 +/- 0.14 | 0.88 +/- 0.14 | 0.88 +/- 0.14 | 0.88 +/- 0.14 | 0.88 +/- 0.14 |
| CytoGLMM | 0.88 +/- 0.14 | 0.88 +/- 0.14 | 0.88 +/- 0.14 | 0.88 +/- 0.14 | 0.88 +/- 0.14 | 0.88 +/- 0.14 | 0.94 +/- 0.12 |
| logRegression | **1 +/- 0** | **1 +/- 0** | **1 +/- 0** | **1 +/- 0** | **1 +/- 0** | **1 +/- 0** | **1 +/- 0** |
| ZAGA | 0.94 +/- 0.12 | 0.94 +/- 0.12 | 0.94 +/- 0.12 | 0.88 +/- 0.14 | 0.94 +/- 0.12 | 0.94 +/- 0.12 | **1 +/- 0** |
| BEZI | 0.88 +/- 0.14 | 0.94 +/- 0.12 | 0.94 +/- 0.12 | 0.81 +/- 0.24 | 0.94 +/- 0.12 | 0.81 +/- 0.12 | 0.38 +/- 0.14 |
| CyEMD | **1 +/- 0** | **1 +/- 0** | **1 +/- 0** | **1 +/- 0** | **1 +/- 0** | **1 +/- 0** | **1 +/- 0** |

**Appendix 0 Table 2.** Specificity of the semi-simulated COVID-19 dataset. The means and standard deviations of the methods are reported across the multiple $\alpha$ values.

| Number of Cells | 1,000 | 2,000 | 5,000 | 10,000 | 15,000 | 20,000 | 4,052,622 |
|---|---|---|---|---|---|---|---|
| diffcyt-DS-limma | 0.96 +/- 0.02 | 0.96 +/- 0.02 | **1 +/- 0** | **1 +/- 0** | **1 +/- 0** | **1 +/- 0** | **1 +/- 0** |
| diffcyt-DS-LMM | 0.73 +/- 0.02 | 0.78 +/- 0 | 0.83 +/- 0 | 0.83 +/- 0 | 0.83 +/- 0 | 0.83 +/- 0 | 0.83 +/- 0 |
| t-test | 0.96 +/- 0.02 | 0.96 +/- 0.02 | **1 +/- 0** | **1 +/- 0** | **1 +/- 0** | **1 +/- 0** | **1 +/- 0** |
| Wilcoxon test | **1 +/- 0** | 0.96 +/- 0.02 | **1 +/- 0** | **1 +/- 0** | **1 +/- 0** | **1 +/- 0** | **1 +/- 0** |
| Kruskal-Wallis test | **1 +/- 0** | **1 +/- 0** | **1 +/- 0** | **1 +/- 0** | **1 +/- 0** | **1 +/- 0** | **1 +/- 0** |
| CytoGLM | 0.96 +/- 0.05 | 0.86 +/- 0.18 | 0.7 +/- 0.23 | 0.66 +/- 0.26 | 0.64 +/- 0.31 | 0.66 +/- 0.28 | 0.57 +/- 0.24 |
| CytoGLMM | 0.91 +/- 0.06 | 0.68 +/- 0.19 | 0.51 +/- 0.31 | 0.47 +/- 0.32 | 0.42 +/- 0.34 | 0.4 +/- 0.35 | 0.43 +/- 0.32 |
| logRegression | **1 +/- 0** | **1 +/- 0** | **1 +/- 0** | **1 +/- 0** | **1 +/- 0** | **1 +/- 0** | 0.96 +/- 0.02 |
| ZAGA | **1 +/- 0** | 0.94 +/- 0 | **1 +/- 0** | **1 +/- 0** | **1 +/- 0** | 0.94 +/- 0 | 0.96 +/- 0.02 |
| BEZI | **1 +/- 0** | **1 +/- 0** | **1 +/- 0** | **1 +/- 0** | **1 +/- 0** | 0.78 +/- 0 | 0.72 +/- 0 |
| CyEMD | **1 +/- 0** | **1 +/- 0** | **1 +/- 0** | **1 +/- 0** | **1 +/- 0** | **1 +/- 0** | **1 +/- 0** |

**Appendix 0 Table 3.** Precision of the semi-simulated COVID-19 dataset. The means and standard deviations of the methods are reported across the multiple $\alpha$ values.

| Number of Cells | 1,000 | 2,000 | 5,000 | 10,000 | 15,000 | 20,000 | 4,052,622 |
|---|---|---|---|---|---|---|---|
| diffcyt-DS-limma | 0.8 +/- 0 | 0.8 +/- 0 | **1 +/- 0** | **1 +/- 0** | **1 +/- 0** | **1 +/- 0** | **1 +/- 0** |
| diffcyt-DS-LMM | 0.36 +/- 0.2 | 0.4 +/- 0.22 | 0.46 +/- 0.26 | 0.46 +/- 0.26 | 0.46 +/- 0.26 | 0.46 +/- 0.26 | 0.46 +/- 0.26 |
| t-test | 0.8 +/- 0 | 0.8 +/- 0 | **1 +/- 0** | **1 +/- 0** | **1 +/- 0** | **1 +/- 0** | **1 +/- 0** |
| Wilcoxon test | **1 +/- 0** | 0.79 +/- 0.03 | **1 +/- 0** | **1 +/- 0** | **1 +/- 0** | **1 +/- 0** | **1 +/- 0** |
| Kruskal-Wallis test | **1 +/- 0** | **1 +/- 0** | **1 +/- 0** | **1 +/- 0** | **1 +/- 0** | **1 +/- 0** | **1 +/- 0** |
| CytoGLM | 0.79 +/- 0.14 | 0.64 +/- 0.29 | 0.37 +/- 0.1 | 0.34 +/- 0.11 | 0.36 +/- 0.17 | 0.36 +/- 0.16 | 0.26 +/- 0.03 |
| CytoGLMM | 0.65 +/- 0.08 | 0.32 +/- 0.02 | 0.25 +/- 0.02 | 0.23 +/- 0.02 | 0.21 +/- 0.03 | 0.21 +/- 0.02 | 0.23 +/- 0.03 |
| logRegression | **1 +/- 0** | **1 +/- 0** | **1 +/- 0** | **1 +/- 0** | **1 +/- 0** | **1 +/- 0** | 0.8 +/- 0 |
| ZAGA | **1 +/- 0** | 0.63 +/- 0.35 | **1 +/- 0** | **1 +/- 0** | **1 +/- 0** | 0.63 +/- 0.35 | 0.8 +/- 0 |
| BEZI | **1 +/- 0** | **1 +/- 0** | **1 +/- 0** | **1 +/- 0** | **1 +/- 0** | 0.36 +/- 0.2 | 0.18 +/- 0.12 |
| CyEMD | **1 +/- 0** | **1 +/- 0** | **1 +/- 0** | **1 +/- 0** | **1 +/- 0** | **1 +/- 0** | **1 +/- 0** |

**Appendix 0 Table 4.** Number of cells per sample of the semi-simulated COVID-19 dataset.

| Patient | Condition | Number of Cells |
|---|---|---|
| CVD001 | base | 176,009 |
| CVD001 | spike | 176,010 |
| CVD003 | base | 244,379 |
| CVD003 | spike | 43,768 |
| CVD006 | base | 201,384 |
| CVD006 | spike | 201,385 |
| CVD007 | base | 199,265 |
| CVD007 | spike | 199,265 |
| CVD010 | base | 191,465 |
| CVD010 | spike | 191,466 |
| CVD011 | base | 189,147 |
| CVD011 | spike | 189,147 |
| CVD013 | base | 197,920 |
| CVD013 | spike | 197,920 |
| CVD019 | base | 266,700 |
| CVD019 | spike | 80,397 |
| CVD012 | base | 154,013 |
| CVD012 | spike | 154,014 |
| CVD020 | base | 198,272 |
| CVD020 | spike | 198,272 |
| CVD023 | base | 201,212 |
| CVD023 | spike | 201,212 |

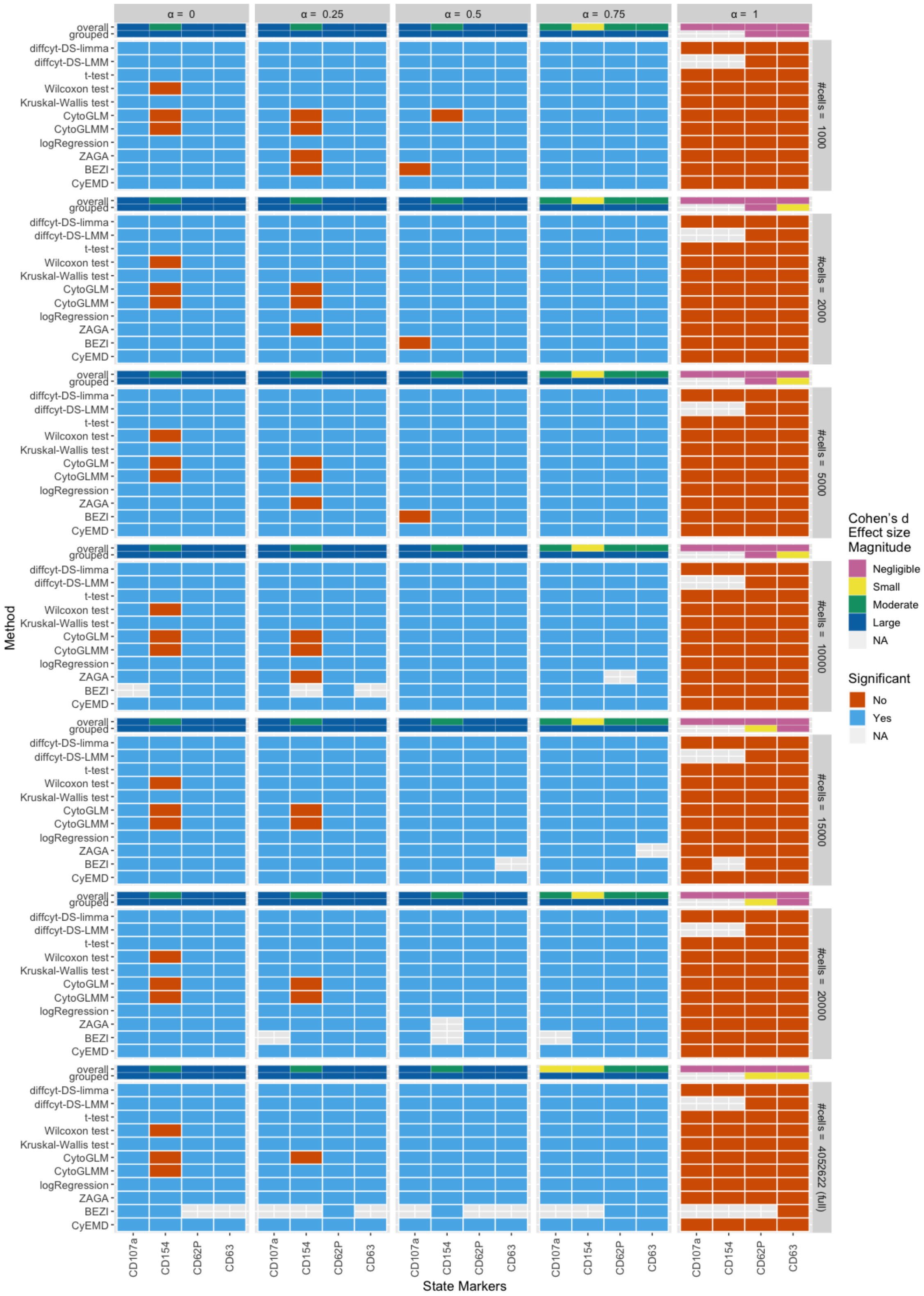


**Appendix 0 Figure 1.** Method results for the state markers of the semi-simulated COVID-19 dataset. The expression of the spiked-in platelets was tested against the baseline samples for differential marker expression. Results colored in blue if the adjusted p-value < 0.05, else in red. Uncolored tiles mean convergence errors of the method for the specific marker. The overall and grouped effect size magnitudes per marker are shown at the top. Overall effect size refers to *Cohen's d* magnitudes using all expression data between two conditions. The magnitudes indicated by grouped effect size are computed in a paired fashion on the median marker expressions per sample. The Wilcoxon test refers to the Wilcoxon signed-rank test.

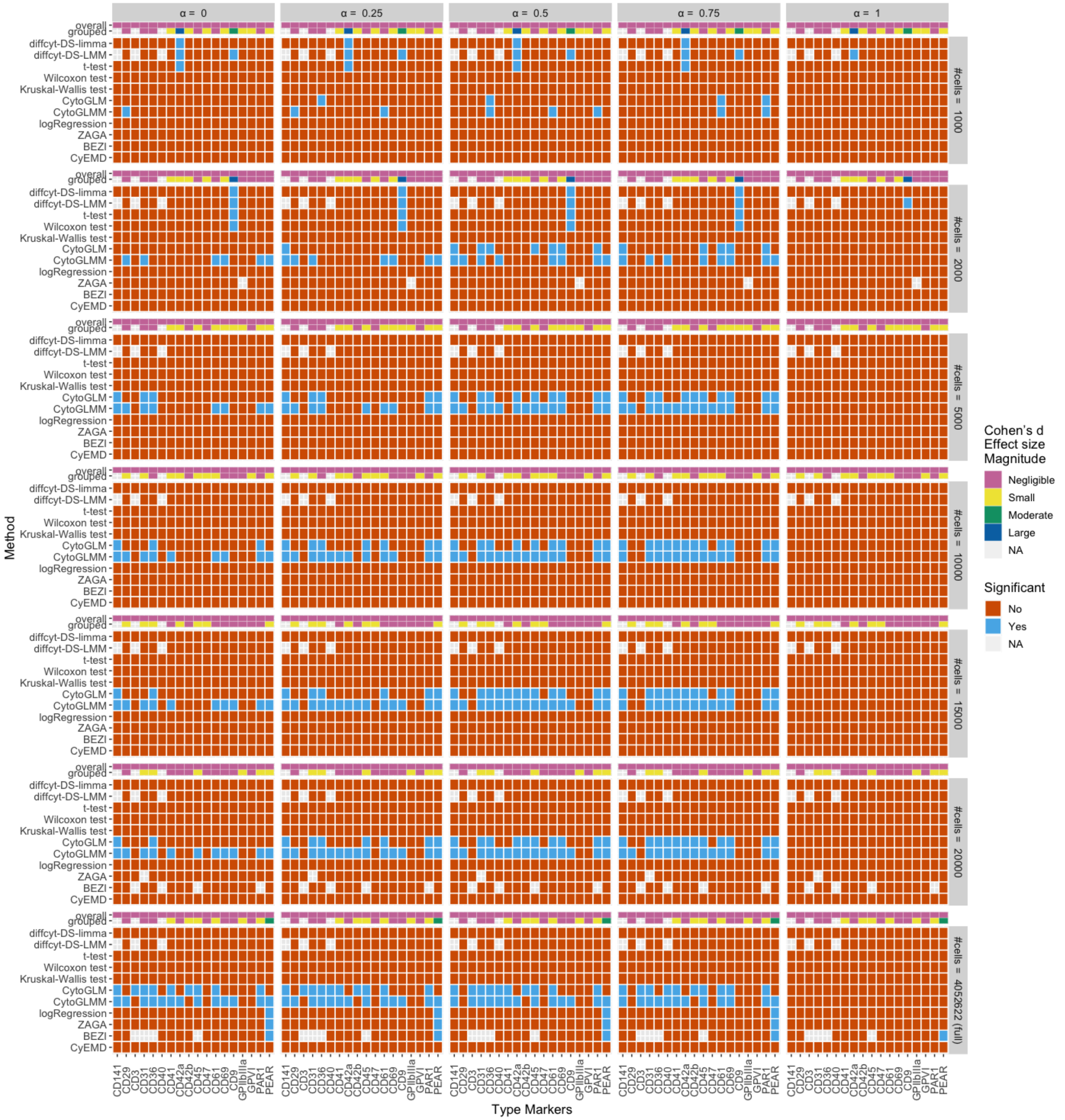


**Appendix 0 Figure 2.** Method results for the type markers of the semi-simulated COVID-19 dataset. The expression of the spiked-in platelets was tested against the baseline samples for differential marker expression. Results colored in blue if the adjusted p-value < 0.05, else in red. Uncolored tiles mean convergence errors of the method for the specific marker. The overall and grouped effect size magnitudes per marker are shown at the top. Overall effect size refers to *Cohen's d* magnitudes using all expression data between two conditions. The magnitudes indicated by grouped effect size are computed in a paired fashion on the median marker expressions per sample. Wilcoxon test refers to the Wilcoxon signed-rank test.

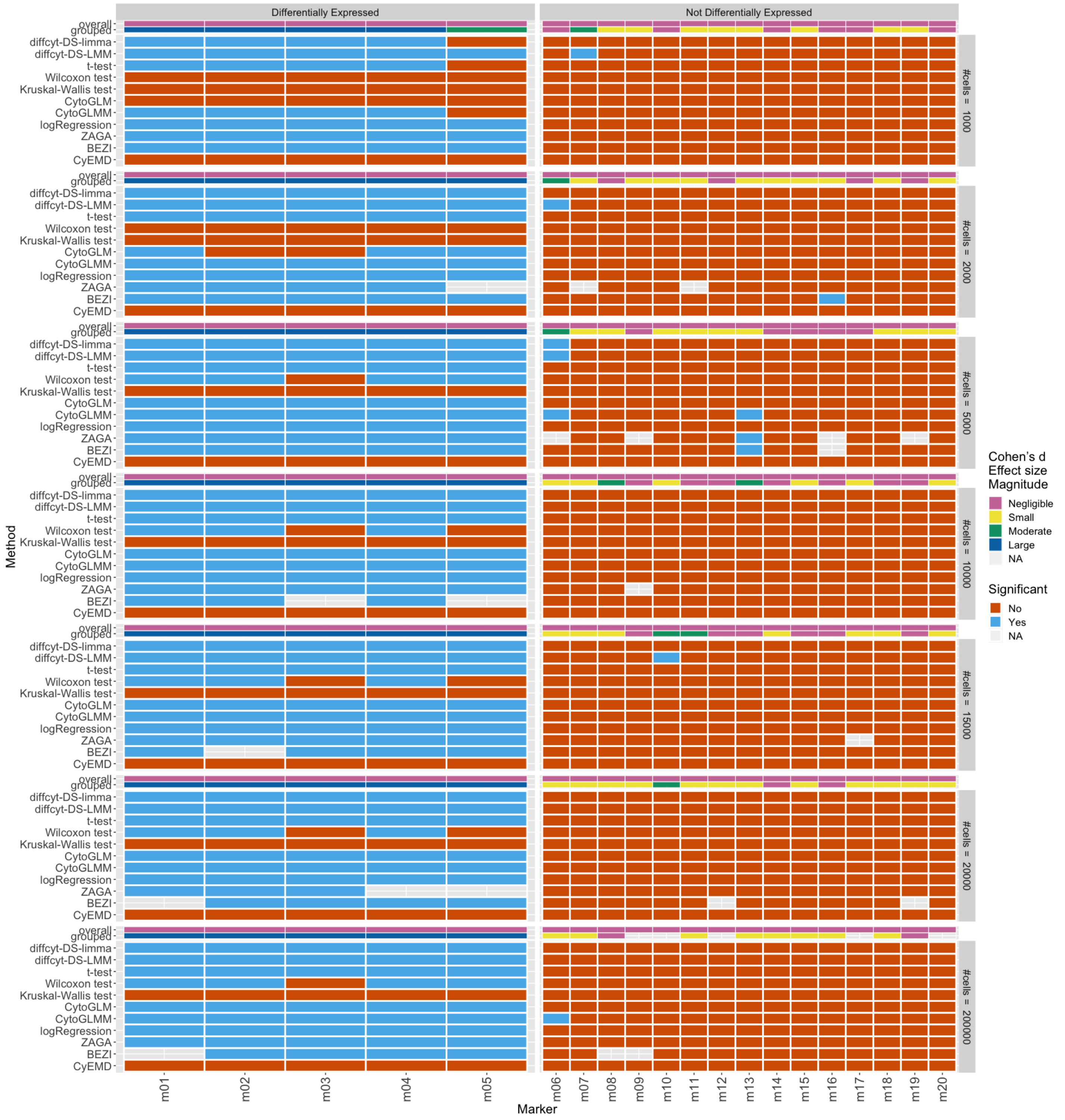


**Appendix 0 Figure 3.** Method results of the simulated dataset from the CytoGLMM package for multiple numbers of cells. The case condition was tested against the control samples for differential marker expression. Results colored in blue if the adjusted p-value < 0.05, else in red. Uncolored tiles mean convergence errors of the method for the specific marker. The overall and grouped effect size magnitudes per marker are shown at the top. Overall effect size refers to *Cohen's d* magnitudes using all expression data between two conditions. The magnitudes indicated by grouped effect size are computed in a paired fashion on the median marker expressions per sample. Wilcoxon test refers to the Wilcoxon signed-rank test.

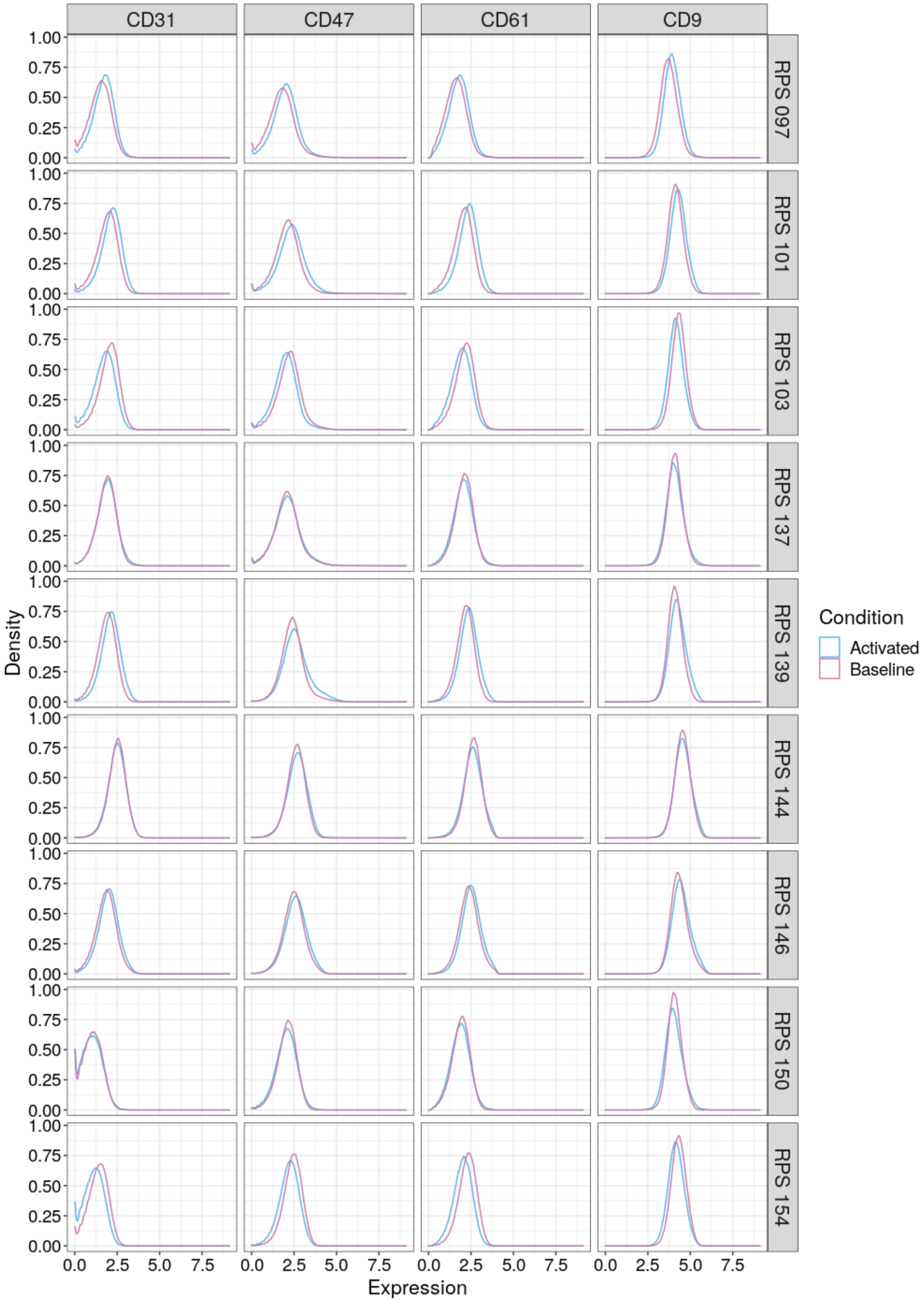


**Appendix 0 Figure 4.** Normalized expression densities for markers with negligible effect size for all patients from the dual dataset.

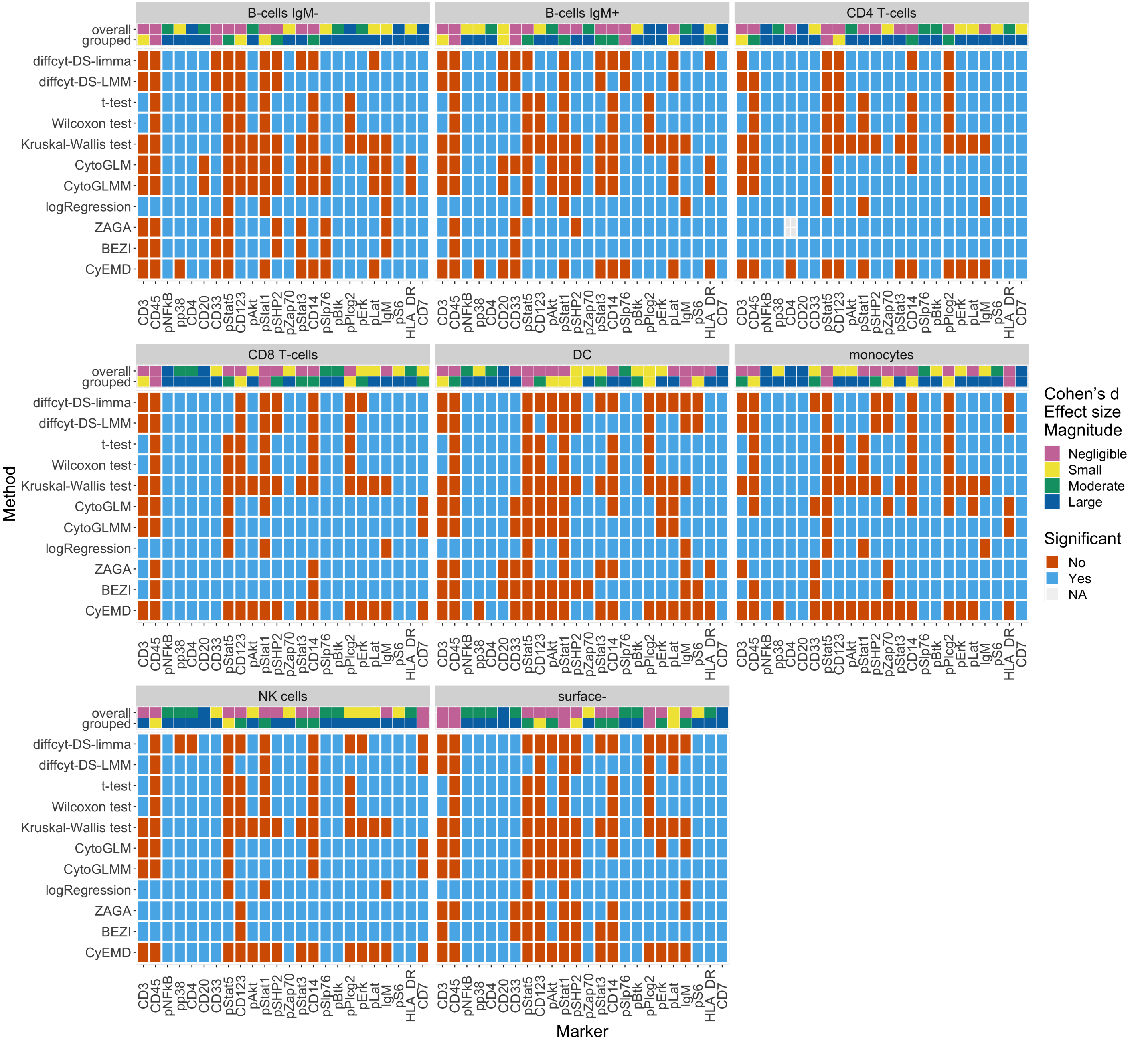


**Appendix 0 Figure 5.** Method results for the PBMC dataset. The BCR-XL condition was tested against the Reference samples for differential marker expression. Tests were performed within each cell type cluster whose annotation was adopted from the CyTOF workflow by ***Nowicka et al.*** (***2019***). Results colored in blue if the adjusted p-value < 0.05, else in red. Uncolored tiles mean convergence errors of the method for the specific marker. The overall and grouped effect size magnitudes per marker are shown at the top. Overall effect size refers to *Cohen's d* magnitudes using all expression data between two conditions. The magnitudes indicated by grouped effect size are computed in a paired fashion on the median marker expressions per sample. Wilcoxon test refers to the Wilcoxon signed-rank test.

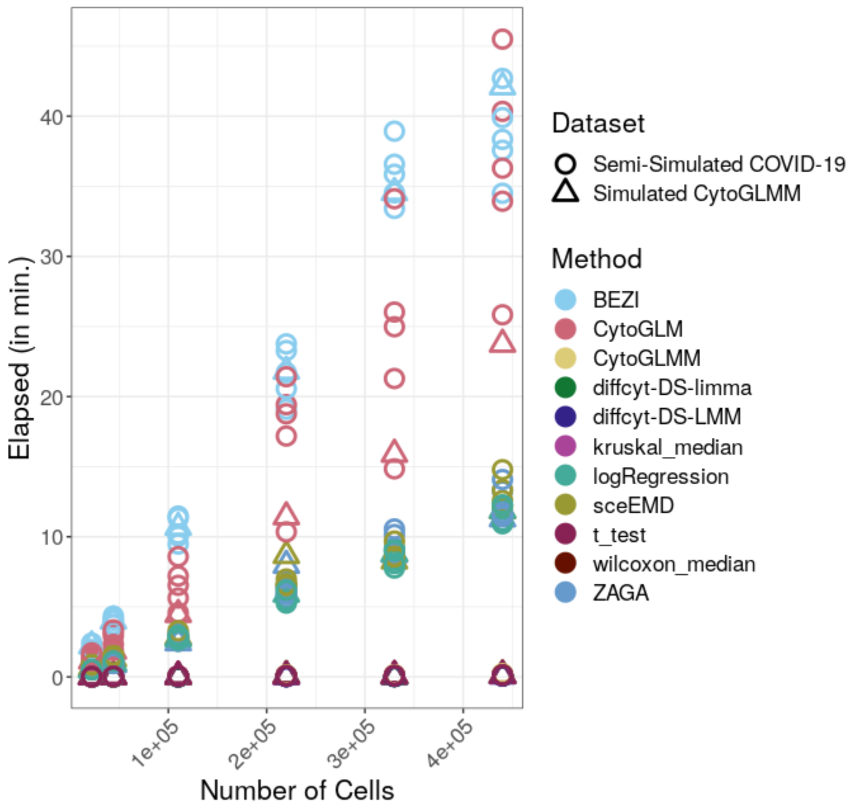


**Appendix 0 Figure 6.** Runtime of the methods on the subsampled datasets, i.e. the semi-simulated COVID-19 datasets and the CytoGLMM datasets. The diffcyt methods and the statistical test are very fast regardless of the sample size. The logistic regression, CyEMD,and ZAGA have moderate runtimes while BEZI and ZAGA are the slowest methods. For all of the methods, the runtime rises linearly with increasing sample size.

**Appendix 0 Table 5.** Number of cells per sample of the dual platelets dataset.

| Patient | Condition | Number of Cells |
|---|---|---|
| RPS 097 | activated | 195,753 |
| RPS 097 | baseline | 142,628 |
| RPS 101 | activated | 303,254 |
| RPS 101 | baseline | 285,330 |
| RPS 103 | activated | 262,761 |
| RPS 103 | baseline | 307,334 |
| RPS 137 | activated | 146,069 |
| RPS 137 | baseline | 229,449 |
| RPS 139 | activated | 234,206 |
| RPS 139 | baseline | 253,639 |
| RPS 144 | activated | 236,872 |
| RPS 144 | baseline | 287,429 |
| RPS 146 | activated | 302,926 |
| RPS 146 | baseline | 295,165 |
| RPS 150 | activated | 161,482 |
| RPS 150 | baseline | 279,348 |
| RPS 154 | activated | 257,038 |
| RPS 154 | baseline | 310,821 |

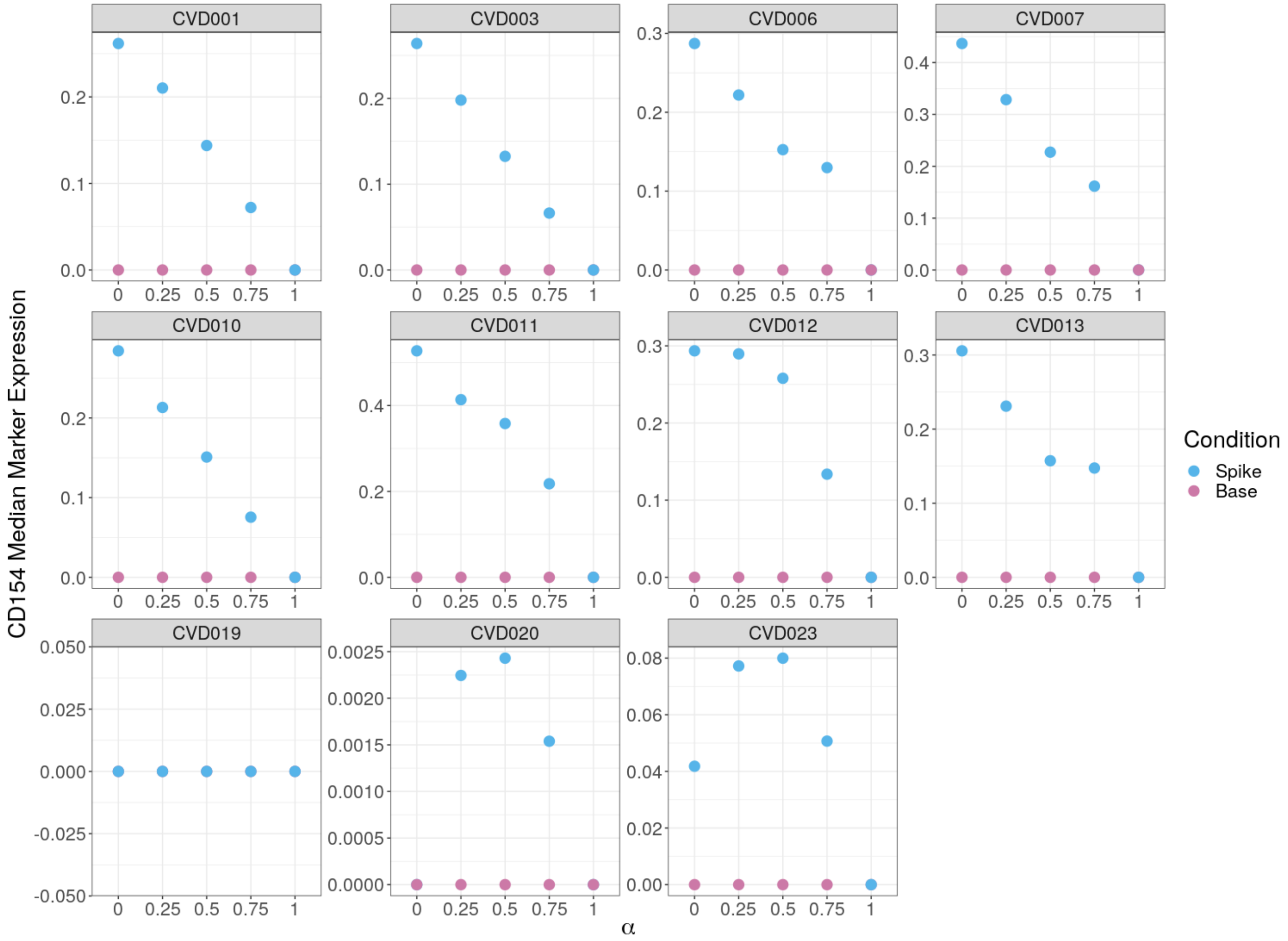


**Appendix 0 Figure 7.** CD154 medians per patient for the full number of cells of the semi-simulated COVID-19 dataset. Medians of the spike condition are higher for $\alpha$=0.25, 0.5, and 0.75 than for $\alpha$=0 for two patients. Additionally, the medians are extremely close to zero.

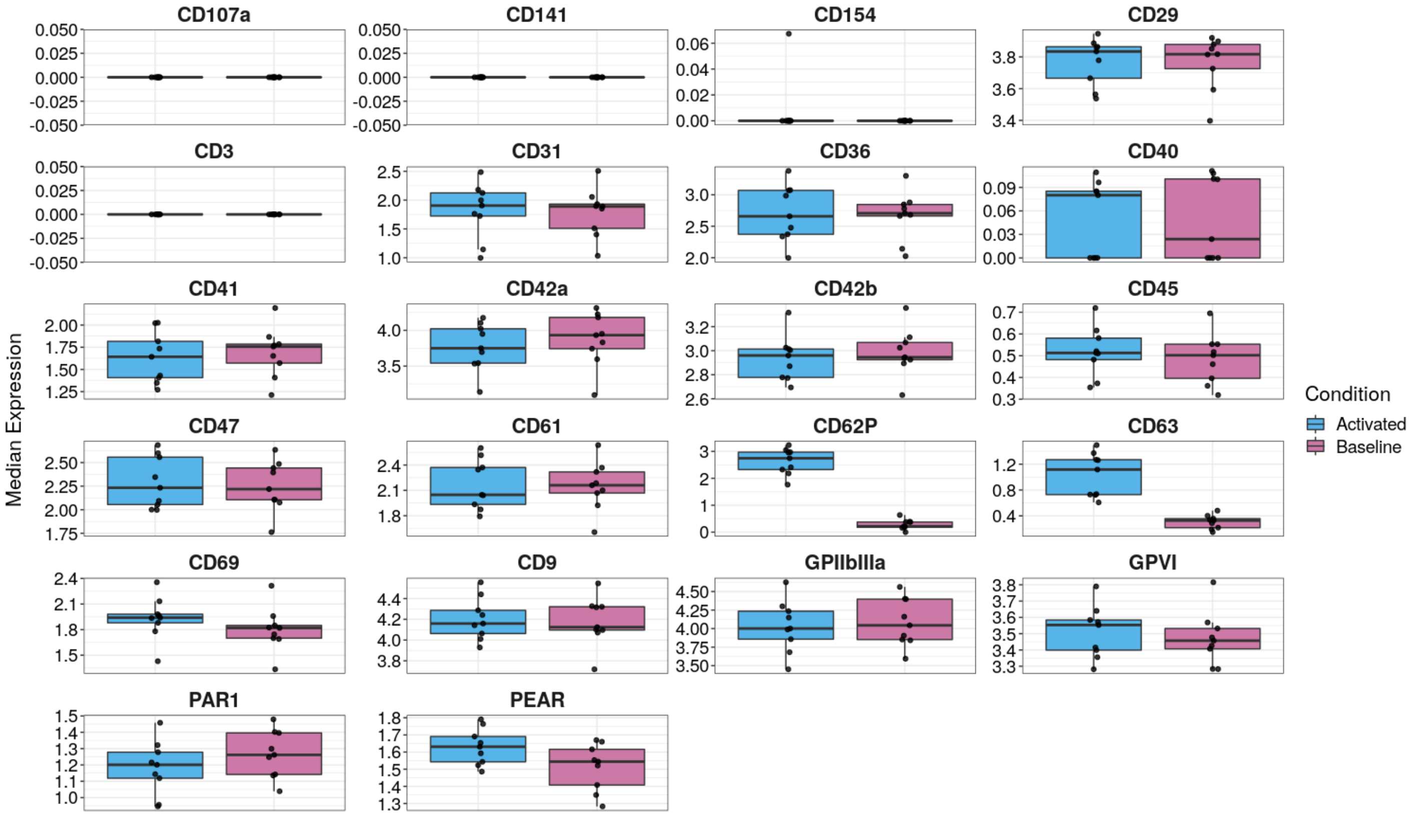


**Appendix 0 Figure 8.** Median marker expressions of all markers from the dual dataset.

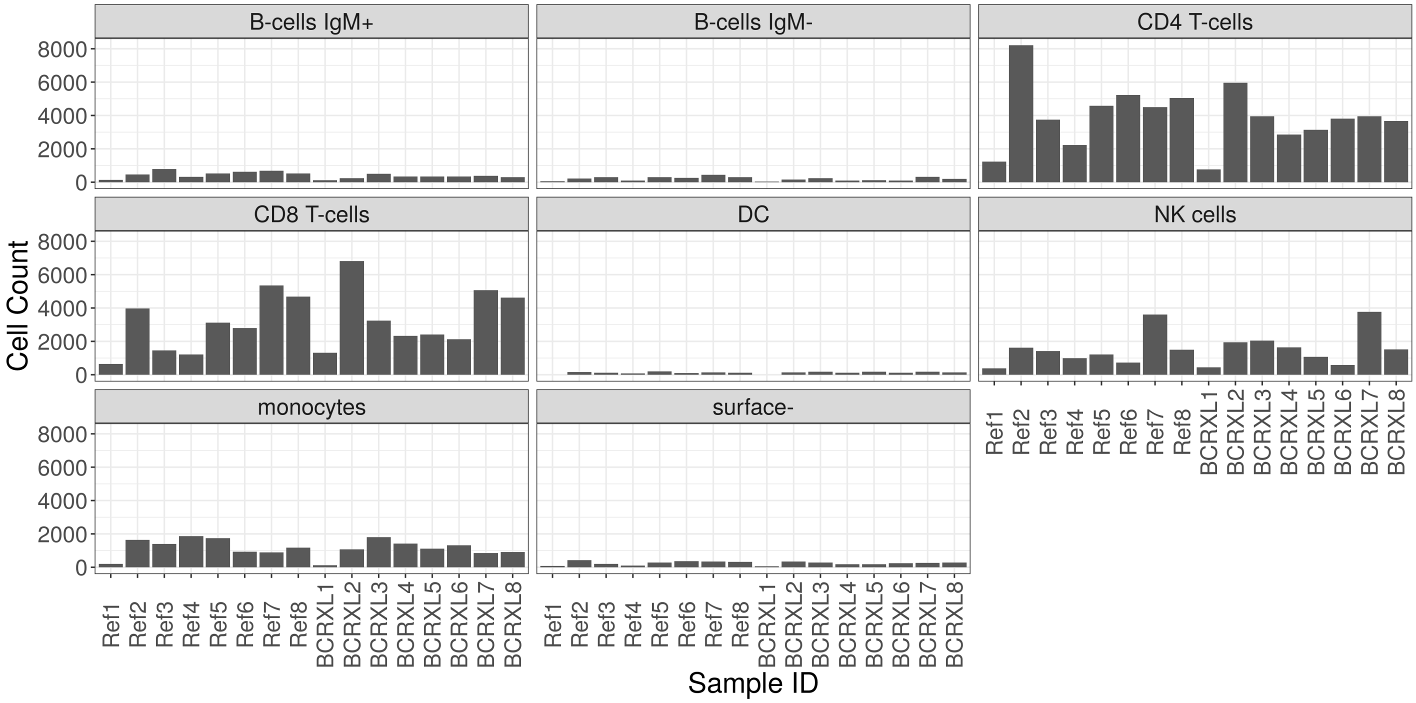


**Appendix 0 Figure 9.** Number of cells per sample and per cluster. CD4- and CD8-T-cells have a much higher cell count for mostly all samples. B-cells, DC, and surface- only have very few cells.

**Appendix 0 Table 6.** Panel of the dual platelet dataset.

| Antigen | Marker Class | Common Name | Biological Function |
|---|---|---|---|
| CD107a | state | LAMP-1 | Cell adhesion, activation |
| CD141 | none | Thrombomodulin | membrane receptor, binds thrombin |
| CD154 | state | CD40L | Interaction with endothelial cells |
| CD29 | type | Integrin $\beta$1 | Collagen receptor unit |
| CD3 | none | TCR-CD3 complex | Adaptive immune response |
| CD31 | type | PECAM-1 | Cell adhesion |
| CD36 | type | GPIV | Collagen receptor |
| CD40 | none | TNFRSF5 | induction of immunoglobulin secretion |
| CD41 | type | Integrin $\alpha$II | Alpha unit of fibrinogen receptor |
| CD42a | type | GPIX | Von Willebrand factor receptor unit |
| CD42b | type | GPIb$\alpha$ | Von Willebrand factor receptor unit |
| CD45 | none | PTPRC | positive regulator of T-cell coactivation |
| CD47 | type | MER6 | adhesion receptor for THBS1 on platelets |
| CD61 | type | Integrin $\beta$3 | Beta unit of fibrinogen receptor |
| CD62P | state | P-Selectin | Cell adhesion, activation marker |
| CD63 | state | LAMP-3 | Cell adhesion, activation marker |
| CD69 | type | CLEC2C | lymphocyte signal transmission |
| CD9 | type | Tetraspanin | Cell adhesion |
| GPVI | type | GPVI | Collagen receptor |
| GPIIbIIIa | type | activated $\alpha$IIb$\beta$3 | Fibrinogen/von Willebrand receptor |
| PAR1 | type | Par1, F2R | Thrombin receptor |
| PEAR | type | JEDI | Platelet Endothelial Aggregation Receptor |

**Appendix 0 Table 7.** Number of cells per sample of the PBMC dataset.

| Patient | Condition | Number of Cells |
|---|---|---|
| Patient1 | BCRXL | 2,838 |
| Patient1 | Ref | 2,739 |
| Patient2 | BCRXL | 16,675 |
| Patient2 | Ref | 16,725 |
| Patient3 | BCRXL | 12,252 |
| Patient3 | Ref | 9,434 |
| Patient4 | BCRXL | 8,990 |
| Patient4 | Ref | 6,906 |
| Patient5 | BCRXL | 8,543 |
| Patient5 | Ref | 11,962 |
| Patient6 | BCRXL | 8,622 |
| Patient6 | Ref | 11,038 |
| Patient7 | BCRXL | 14,770 |
| Patient7 | Ref | 15,974 |
| Patient8 | BCRXL | 11,653 |
| Patient8 | Ref | 13,670 |
